# MortX: A Domain Generalization Benchmark for Mouse Cortex Segmentation and Registration

**DOI:** 10.1101/2024.11.30.626208

**Authors:** Asim Iqbal, Romesa Khan, Edith M. Schneider Gasser, Theofanis Karayannis

**Author notes:** Corresponding authors: Asim Iqbal and Theofanis Karayannis.

## Abstract

Mesoscale understanding of human brain development is crucial for understanding neurodevelopmental disorders. By applying AI techniques to analyze high-resolution, multi-modal brain imaging datasets across postnatal ages, researchers can study cortical development at the granular level. We introduce MortX, a benchmark dataset of the developing mouse cortex that captures multiple postnatal stages with annotations for distinct anatomical and functional subregions and layers. MortX features high-resolution imaging data including bright-field and fluorescence-labeled neuronal markers. We developed a standardized cortical atlas of genetic markers and manually registered it to brain section images for ground-truth labeling. The dataset serves as a benchmark for domain generalization in neuroimaging, enabling both classical and deep learning models to be trained on source brains and tested on unseen targets. Our results demonstrate generalized model performance and structural invariance across ages. We open-source MortX as a community resource for mouse brain segmentation and registration, emphasizing domain adaptation. This dataset addresses key challenges in mouse brain imaging and advances machine learning models that will help unravel neurodevelopmental disorders.

## 1 Introduction

The mammalian cortex forms the complex substrate for most higher-level cognitive processes and is organized into histologically and functionally distinct sub-regions. Radially, it forms a laminated structure, comprising layers distinguished in cytoarchitecture, neuronal morphology, and gene expression. In the tangential plane, the layered cortex is further divided into functionally specialized areas, each defined by connectivity to other brain areas, cytoarchitecture, and molecular profiles (O’Leary et al. [2007]). Neuroscientists are gathering increasing information, and charting changes in neuronal connectivity and functional activity patterns through the course of mammalian brain development. However, to navigate any such high-dimensional anatomical or functional data, this information needs to be precisely mapped to a specific cortical subregion. This step critically depends on the unambiguous anatomical delineation of the cortex, and therefore, on the availability of a sufficiently detailed anatomical reference atlas of the mammalian cortex. In this regard, numerous efforts among the neuroscience community have been directed toward fine-structure annotation of the adult mouse brain cortex (Atlas [2011], Jin et al. [2022]), the most widely studied mammalian model in neuroscience. However, fine annotation of the mouse cortex at earlier developmental time points has remained an uncharted territory (Kronman et al. [2024]). This precludes a deeper investigation of the anatomical and functional organization of the developing cortex, as well as identifying the cortical underpinnings of behavioral alterations during the development. To address the imperative need among the developmental neuroscience community, to access a detailed cortical reference atlas for the postnatal mouse brain, we have created a set of cortical reference atlases across salient developmental time points, namely: P4 and P14, with fine-structure annotation of cortical subregions, based on differential gene expression of layer-specific and region-specific neuronal markers (Molyneaux et al. [2007]). Moreover, we have modified the already existing Allen P56 mouse brain atlas, to create an integrated series of developing mouse cortical reference atlases, which can collectively provide a valuable neuroinformatics tool. Most importantly, the annotation of unique cortical subregions allows this reference atlas series to be directly utilized to create labeled ground-truth data for building robust and fully automated deep learning-based approaches for cortical segmentation and atlas registration (Iqbal et al. [2019], Ronneberger et al. [2015]).

Despite the impressive capabilities demonstrated by the above-mentioned deep neural networks when applied to identically distributed data, they can encounter significant difficulties when faced with shifts in the data distribution. Domain generalization aims to address the performance degradation of the trained models when confronted with new, previously “unseen” domains during inference, without the need for explicit domain-specific training or adaptation (Wang et al. [2021], Zhou et al. [2023]). However, currently, there is a scarcity of domain generalization datasets in neuroimaging. In particular, despite the extensive use of mice as a mammalian model in neuroscience research, there is no existing benchmark dataset that addresses the domain generalization issue in the mouse brain imaging community. To facilitate the analysis of the diverse and large-scale mouse brain imaging data, we introduce **MortX** (**Mo**use co**rt**e**X**), a domain generalization benchmark dataset that incorporates mouse brain images labeled with multiple genetic markers and acquired through various imaging modalities, across multiple mouse developmental time points. Our dataset will not only be useful for the community to map the developing mouse brain images to a generalized atlas but also to build domain-invariant models for mouse cortex segmentation and atlas registration-based tasks.

## 2 Background and Related Work

### 2.1 Mapping the developing mouse cortex

The Allen Mouse Brain CCF (Atlas [2011]) is an outstanding initiative in this regard, which provides comprehensive annotation of cortical subregions. However, standardized atlases with fine annotations of the mouse cortex at earlier developmental time points do not exist. Furthermore, even in the case of the adult mouse cortex where such an atlas is available, it is only registered to a specific anatomical template (e.g. a Nissl-labelled P56 mouse brain in the case of the two-dimensional (2D) Allen adult mouse atlas). It needs to be registered to the target brain. Notably, the registration of the reference atlas onto the experimental brain data is a non-trivial problem due to substantial genetic variability across different animal brains and untrackable domain shifts caused by the multiple steps before and during image acquisition. Classical approaches to the problem of the atlas image registration constitute applications of affine and non-rigid transformations to the reference coordinate system, to align the tissue boundaries within the input brain image (Klein et al. [2010], Kutten et al. [2017]).

However, despite being largely semi-supervised and dependent on user-defined parameters, these registration algorithms show limited performance in delineating individual regions. Geometrically precise boundary handling becomes a graver challenge in the case of mapping fine-structural annotations, where the higher number of ontological subregions and their compact arrangement complicate the accurate assignment of labels. Recently, however, deep convolutional neural networks have been successfully applied to fully automate the task of brain reference atlas registration (Iqbal et al. [2019], Shakeri et al. [2016], Mehta et al. [2017], Xiao et al. [2021]). In segmentation-based tasks, each brain region is considered a unique instance of an object, which the deep neural network learns to detect, classify, and segment in an input brain image, through recognition of core anatomical (e.g. tissue morphology- and texture-based) features. The source brain (moving) image section is mapped to the target brain (fixed) image section via spatial transformations using deep learning-based registration techniques for registration-based tasks (Razzaq and Iqbal [2024], Mahmood et al. [[2020], [2023], [2024]a,b], Payette et al. [2021]).

### 2.2 Domain Generalization in brain imaging

While domain adaptation has been a longstanding challenge in the computer vision community, recently, some important domain generalization datasets have emerged with a specialized focus on medical and brain imaging. One prominent dataset is the Alzheimer’s Disease Neuroimaging Initiative (ADNI) dataset (Jack Jr et al. [2008]). ADNI is a large-scale longitudinal study that includes MRI, PET, and other clinical data from individuals with Alzheimer’s disease, mild cognitive impairment, and healthy controls. The dataset provides a valuable resource for developing and evaluating domain generalization methods in the context of neurodegenerative diseases. Another notable dataset is the Multi-Modality Whole Heart Segmentation (MM-WHS) Challenge dataset (Zhuang [2018]). This dataset contains cardiac MRI scans from different institutions and imaging protocols, making it a suitable benchmark for evaluating domain generalization approaches in cardiac image segmentation. Furthermore, the Human Connectome Project (HCP) dataset (Elam et al. [2021]) is a comprehensive resource for studying brain connectivity and function. It includes high-quality structural and functional MRI data collected from a large number of healthy participants. The HCP dataset has been widely utilized in the domain generalization literature to investigate the transferability of models across different scanning sites and acquisition protocols. Roels et al. [2019] highlight another effort for domain adaptive benchmarking of volume electron microscopy data, such as domain adaptation from isotropic FIB/SEM volumes to anisotropic TEM volumes. Another evolving domain adaptation benchmark among the medical imaging community is TissueNet, which encompasses both human histopathology and fluorescence microscopy images (Greenwald et al. [2022]).

While these domain generalization datasets are available for human brain imaging, such datasets are scarce for animal models. In particular, no such benchmark exists to our knowledge, which addresses the problem of domain generalization or domain shift robustness in mouse brain imaging. Mouse is one of the most widely studied mammalian models by the neuroscience community, due to its expedient developmental timeline, during the multiple steps before and during image acquisition, as well as the genotypic and phenotypic homology with humans. To better understand the mouse brain, neuroscientists label the tissue with numerous genetic markers and acquire brain images using a range of imaging modalities and scanner types. This has created canons of large-scale, highly-varied data, acquired using diverse methods of imaging the mouse brain. In order to leverage AI-based models to analyze such data in a robust and high-throughput manner, the distributional variations in this data must be reconciled by domain generalization-based approaches. To address this critical need in the neuroscience community, we present a domain generalization benchmark for the mouse brain, which encompasses multiple genetic markers and diverse imaging modalities, across three animal age groups: P4, P14, and P56.

## 3 Results

### 3.1 Developing cortical atlas generation

#### 3.1.1 Mapping of P56 mouse isocortex onto P14 and P4 isocortices

The adult mouse brain reference atlas from the open-source Allen Brain Map reference atlases was used as a spatial coordinate framework to map cortical subregions. Each image section of the detailed adult mouse brain reference atlas was matched to a corresponding section of the target developing mouse brain reference atlas, at ages P14 and P4, respectively. The corresponding image sections were selected based on the position of the sagittal plane along its lateromedial axis, determined by the appearance of the landmark structures. In the first stage, a scalar vector graphics (SVG) file of a P56 reference brain section was overlaid onto the corresponding P4/P14 reference brain section image. Only the P56 reference cortex was retained and all the other brain structures were removed. The external boundary of the P56 reference cortex was completely registered to the cortical boundary of the underlying developing mouse reference section. This is executed through point-by-point adjustment of individual vectors defining the cortical sub-region boundaries. This results in minimal alteration of the original shape and relative sizes of the cortical sub-regions. The registration process is aimed towards an elliptical approximation of the brain’s cortical surface, and region-wise boundaries that observe the topological notion of simply connected spaces (Krantz et al. [1999]). Each cortical sub-region of the cortex, e.g., layer 2/3 of the visual areas, can be modeled as a conical frustum with an ellipsoidal head of Gaussian curvature, *K*, given in Cartesian coordinates *x*,*y* and *z*:

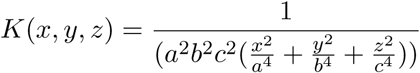

where the semi-axes of the ellipsoid are of lengths *a*, *b*, and *c*. All layers within the same functional region can be modeled to follow the same curvature. The edge of each conical frustum can be described by:

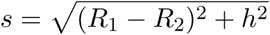

where *s* is the slant height, *h* is the perpendicular height, *R*_1_ is the radius of the superficial cortical layer and *R*_2_ is the radius of the deeper cortical layer.

More specifically, this process of registration is geared towards achieving a conformal mapping (Krantz et al. [1999]) between a given P56 reference brain section and the corresponding P4/P14 reference brain section image. The goal of image registration is to find a transformation that most optimally maps the spatial features in one image to the ones in the target image. Conformal mapping is a mathematical transformation that preserves local angles and the shape of objects in an image. The motivation for considering conformal mapping is rooted in seminal work from the biological field of morphometrics, which has proposed that the natural process of growth produces conformal deformations in an organism’s body (Thompson [1992], Petukhov [1989], Milnor [2010], Tufail [2017]). Based on this historical evidence, we argue that the growth of individual regions in the mouse brain across postnatal developmental timepoints, such as P4, P14, and P56, must be captured by a conformal transformation. This transformation will belong to the Mobius subgroup of conformal transformations, which preserves cross-ratios between the source and target images, i.e. corresponding young and adult mouse brain sections. Specifically, a cross-ratio of four points *A, B, C, D* ∈ C is defined as following:

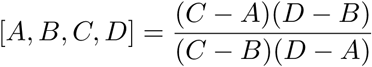

where *A, B, C, D* represent the invariant landmark points on source and target images. In our particular case, these landmark points lie on the boundaries of the distinct brain regions, in both young and adult mice. C denotes a complex plane. This transformation should minimize angular distortion and preserve the structural shape of brain regions in the P4/P14 mouse brain section that is registered to the corresponding adult (P56) mouse brain section image.

### 3.2 Cortical gene expression-dependent arealization and layerization

Anatomical boundaries were delineated for 13 functionally distinguished cortical areas, namely: insula (gustatory and visceral areas), somatomotor areas, somatosensory area, posterior parietal association areas, visual areas, retrosplenial areas, ectorhinal areas, entorhinal areas, frontal pole, orbital area, anterior cingulate area, prelimbic area, and infralimbic area. Laminar boundaries were further specified according to the layerization within each cortical region, ranging between 1 to 6 layers: layer 1, layer 2/3, layer 4, layer 5, layer 5/6, and layer 6. In-situ hybridized (ISH) brains labeled with the selected markers, were obtained from the open-source Allen Developing Mouse Brain Atlas database (Atlas [2011]). The P56 cortex registered to the developing mouse brain reference section after the first stage of registration (referred to as the *pre-registered cortex* section hereon) was overlaid with the image of the ISH brain section. This ISH section is labeled with the neuronal marker that is previously established to differentially label a specific cortical area(s) or layer(s) at the particular developmental age (P4/P14), therefore, outlining the boundaries of this cortical area(s) or layer(s).

In the second stage, the ISH brain section image was registered to the pre-registered cortex section by manually applying a form of non-rigid transformation that maximizes alignment of the external cortical boundary of the ISH brain image with the external cortical boundary of the underlying pre-registered cortex section, while the visually minimizing surface distance between the whole brain boundaries of target and template section. This process of coarse registration of the ISH brain image is described in the Methods section. In the third stage, the internal boundaries of cortical subregions are refined with reference to genetic marker expression in the registered ISH brain image, by point-by-point vector adjustment within the pre-registered cortex section. The process for fine-registration of cortical subregions is also illustrated in the Appendix. In addition to the ISH brain images for the selected neuronal markers, their corresponding gene expression heat maps were also used to aid the localization of the structure-specific gene expression. The genetic neuronal markers for the developing brain that were utilized to localize cortical subregion boundaries (cortical arealization and cortical layerization) are described below:

#### 3.2.1 P4 arealization and layerization

We utilized transcription factors Bhlhe22 (Basic Helix-Loop-Helix Family Member E22), Rorb (RAR-related orphan receptor beta), Lmo4 (LIM Domain Transcription Factor LMO4) and cell adhesion molecules, Cdh6 (Cadherin-6) and Cdh8 (Cadherin-8), to mark the areal boundaries between cortical subregions. Bhlhe22, Rorb, and Cdh6 are differentially expressed across cortical laminae, also making these appropriate indicators for mapping layer-wise boundaries.

#### 3.2.2 P14 arealization and layerization

Rorb marker expression is regulated at cortical layer and areal borders, highlighting the separation between these areas. Bhlhe22, Bcl6 (B-cell lymphoma 6), Lmo8 and Cdh8 are additionally informative in demarcating cortical subregion-wise boundaries.

### 3.3 Fine-structure color annotation of developing cortical subregions

After the genetic label-guided refinement of cortical subregion boundaries, each cortical subregion was assigned a unique RGB color code. Anatomical sub-divisions of any cortical area, into dorsal and ventral regions, were united under an identical color label. Laminar subdivisions of layer 6, namely: layers 6a and 6b, were also merged. The visceral and gustatory areas were united into the insular cortex, according to the previous version of the Allen adult mouse brain atlas. **Figure 1** also demonstrates color labels for the corresponding atlas images where **Figure 2** shows selected reference atlas sections mapped to their corresponding brain sections.

**Figure 1:**
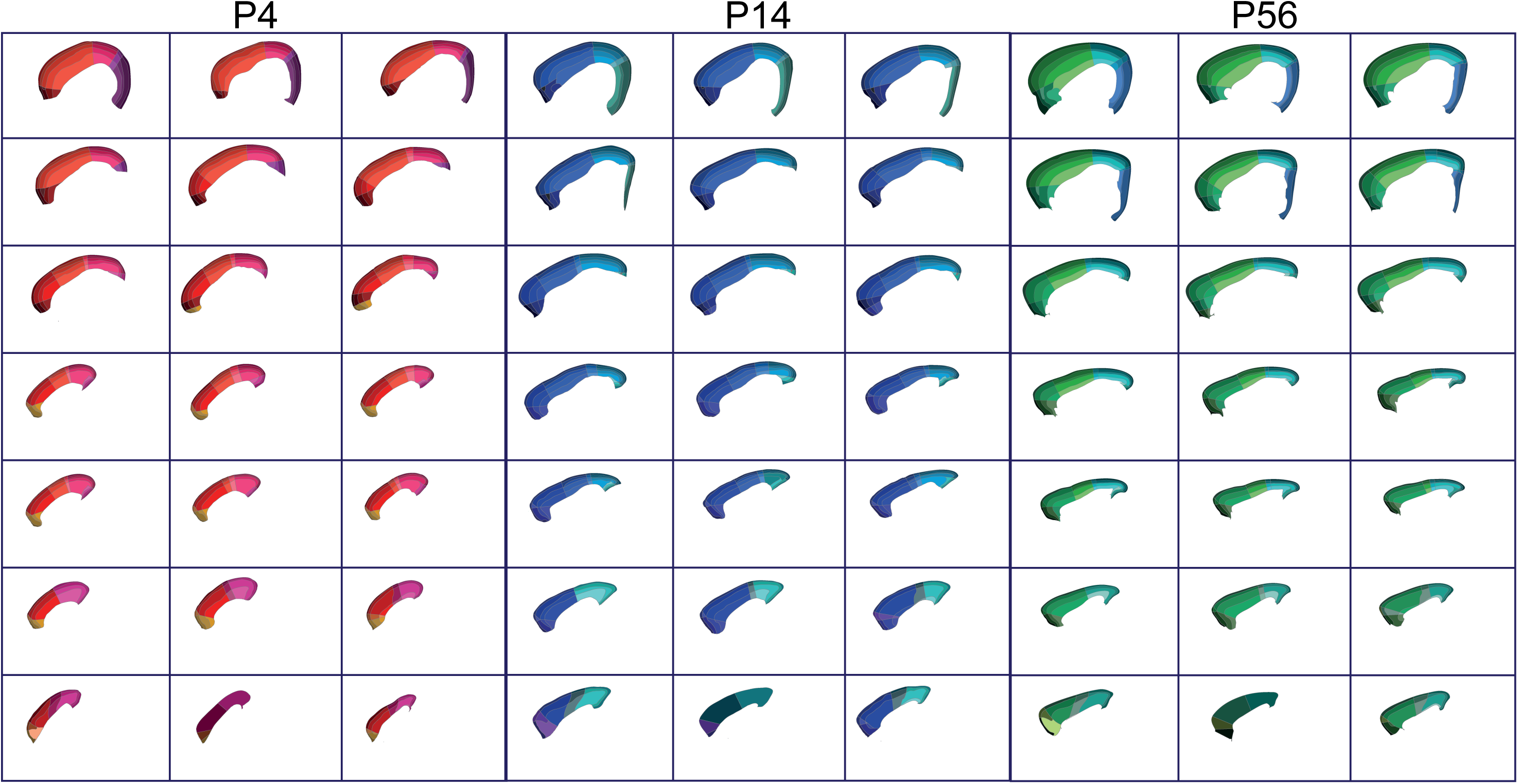
High-level overview of the developing cortical atlas across salient mouse postnatal ages. The mapping of each atlas section is represented in the same order across development.

**Figure 2:**
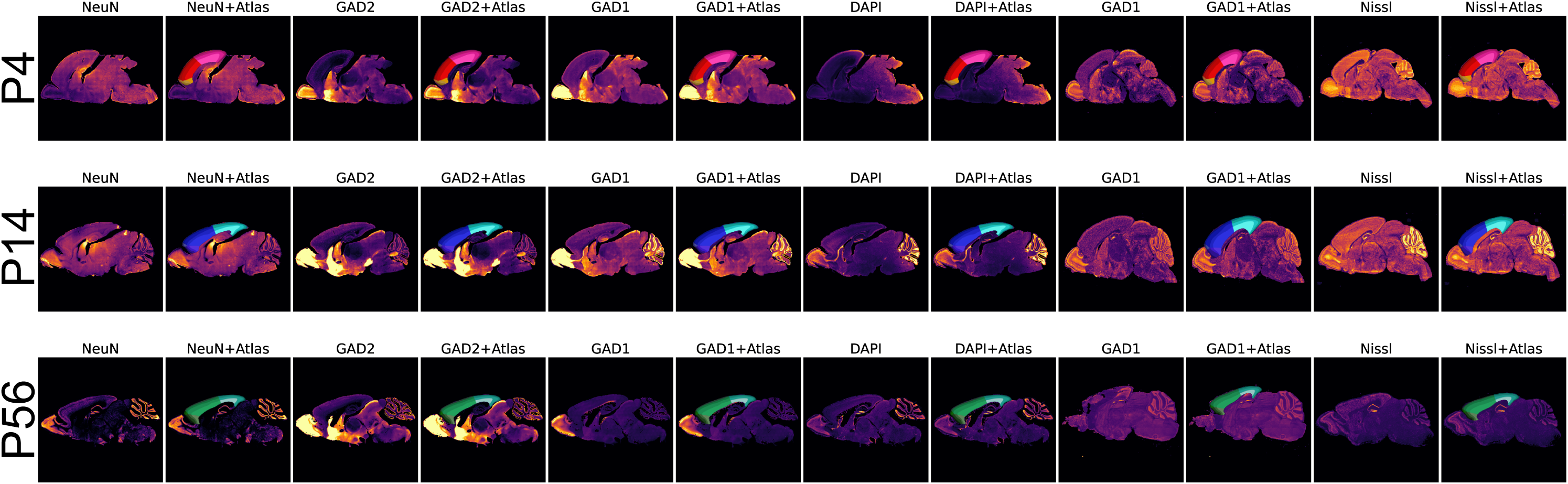
The structural and domain variability of the dataset is represented using a few samples of mouse brain sections, covering different genetic markers with their overlaying cortical atlas labels.

### 3.4 Domain generalization dataset of developing mouse cortex

We introduce a novel domain generalization benchmark for the developing mouse brain cortex (MortX). This dataset incorporates the domain distribution shift at three levels:

**Structural variation**: The acquired brains comprise of three mouse postnatal age-groups: P3-P4, P14, P56-P60, here onward referred to as P4, P14 and P56 for simplicity. This data spans changes in brain tissue morphology across the developmental timeline. Each brain section image is provided as a high-resolution grayscale image. **Figure 2** demonstrates the diversity and richness of our domain generalization benchmark. In this figure, representative lateral and medial brain sections across all mouse postnatal ages are presented, exemplifying the cortical structural variation that is found within and across different mouse developmental time points. Moreover, the domain variation across commonly used brain tissue labeling techniques covered in this benchmark, including genetic markers and nuclear stains, are presented therein. Each panel qualitatively displays the characteristic image attributes, such as luminance, intensity, contrast, and unique spatial expression patterns, associated with all the brain markers included in MortX.

**Imaging modality**: The mouse brains were acquired using two different imaging techniques: bright-field microscopy (BFM) and fluorescence microscopy (FM).

**Tissue labeling**: The mouse brains were labelled using stains and genetic markers that are most widely used in the brain: For nucleic acid stains, Nissl is traditionally used to label all cells in the brain by staining their rough endoplasmic reticulum (Kádár et al. [2009]); DAPI is a DNA-binding fluorescent stain commonly used for nuclear quantitation of nervous tissue(Tarnowski et al. [1991]). NeuN (pan-neuronal marker) is widely used to label the nuclei and perikarya of most neuronal cell types (Gusel’Nikova and Korzhevskiy [2015]). For excitatory neurons, Camk2a is a widely used synaptic marker of excitatory neuron populations in the mammalian central nervous system (Wang et al. [2013]). In addition, conventional inhibitory neuronal markers, GAD1 and GAD2, label overlapping populations of GABAergic neurons (Trifonov et al. [2014]).

**Ground-truth labeling**: The ground-truth cortical region-wise segmentation labels were generated by manual registration of the developing mouse cortical atlas onto each mouse brain in the MortX dataset. Specifically, each 2D brain section image was overlaid by the corresponding reference section in the mouse age-specific cortical atlas, where each brain region was assigned a unique color code. Human experts manually registered each 2D brain section with scalar vector graphics files of the reference atlas in a scalar vector graphics editing software.

### 3.5 Domain generalization benchmarking tasks for developing mouse cortex

Each parent task comprises two sub-tasks, on which the models are evaluated:

**Whole-cortex segmentation**: This sub-task is designed to measure any performance degradation of deep-learning-based segmentation models in segmenting the complete mouse brain cortex, under an age-dependent brain structural distribution shift and/or a brain marker-induced domain shift.

**Cortical atlas registration**: This sub-task aims to evaluate the performance of both classical and deep-learning-based approaches for mouse cortical atlas registration, in the context of age-related variation in the cortical boundary and/or variance caused by the tissue labeling technique.

#### 3.5.1 Domain generalization tasks’ complexity

Neuroscientists label mouse brain tissue with a large variety of genetic markers and cellular stains, depending on their scientific question of interest. Each such staining or genetic label represents a unique domain or distributional shift in the brain imaging data. This complexity and diversity necessitate that the deep learning-based techniques employed for analyzing neuroimaging data analysis are resilient to these domain shifts. This robustness is crucial to ensure the accuracy and reliability of neuroimaging data interpretation. To answer this need, we pose a domain generalization task where the source and target domains comprise mouse brain images that are labeled for diverse genetic markers and nuclear stains, within a single mouse age group. Specifically, we present this task at multiple levels of complexity, for instance:

**Simple**: The model is trained on 4 source domains (brain markers) and tested on 1 unseen target domain. For example, a model is trained on P3-P4 brains labeled for GAD1, GAD2, NeuN, and Nissl, and tested on DAPI.

**Intermediate**: The model is trained on 2 source domains and tested on 3 unseen target domains, where the source and target domain sets have distinguishable target cell populations. For example, a model is trained on P3-P4 brains labeled for the inhibitory markers GAD1 and GAD2 and tested on pan-neuronal and pan-cellular markers, NeuN, Nissl, and DAPI.

**Complex**: The model is trained on 1 source domain and tested on 3 unseen target domains, where the source and target domain sets have different expression patterns. For example, a model is trained on P3-P4 brains labeled for the inhibitory markers GAD1, and tested on pan-neuronal and pan-cellular markers, NeuN, Nissl, and DAPI. For a complete range of all possible combinations of source and target domain sets, please look at **Figure 3**.

**Figure 3:**
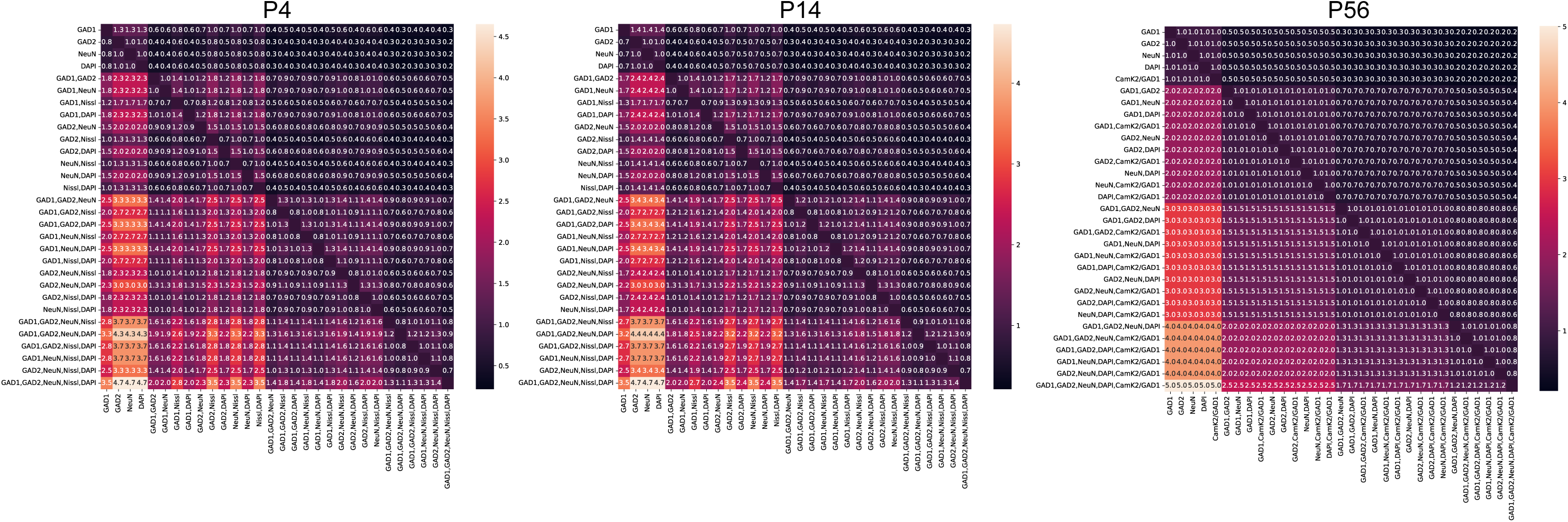
The possible combinations for training on the source (rows) and testing on unseen target (columns) domains are presented with their train/test ratio numbers on top. The color bar shows the ratio of data size (train/test). The dark colors represent tasks with higher levels of complexity, whereas light colors depict the easier tasks with include a large training set for domain robustness.

#### 3.5.2 Multi-level domain generalization tasks

While tasks 1 and 2 can be considered domain generalization problems of intermediate complexity, we also introduce a domain generalization task of advanced complexity, where the source and target domains are separated by a two-level domain shift of animal age as well as tissue labeling technique. e.g. a model trained on P4 brains labeled for GAD1 will be tested on P56 brains labeled for NeuN. For anatomical and functional connectivity analyses of the developing cortex, it is imperative to map brain images from all animal ages onto a detailed cortical atlas, which is partitioned into distinct functional areas and laminae. Therefore, the segmentation and registration models utilized for such analyses must also be able to map the cortex at the cortical sub-region level. Notably, the developing mouse cortical atlas can be leveraged to evaluate all model performances on tasks 1-3 at a more granular level of cortical functional regions as well as individual cortical layers, thus providing the community with a much-needed domain generalization benchmark for fine-scale segmentation and registration tasks.

### 3.6 Benchmarking experiments

#### 3.6.1 Domain generalization across cortical developmental timepoints

**Models**: In this sub-task, we perform segmentation of the complete mouse cortex on brain images, labeled with a pan-neuronal brain marker (NeuN). To evaluate cross-age structural invariance, we train each model on three different dataset splits, detailed under rows 1-3 in **Table 1**. We benchmark two supervised deep-learning-based segmentation models, SeBRe (Iqbal et al. [2019]) and 2D U-Net (Ronneberger et al. [2015]), which are commonly used by the neuroimaging community. Additionally, we benchmark two model variants of a classical registration model, ANTS (Avants et al. [2011]): Affine and Affine+Non-Rigid. The experimental design is detailed under rows 1-3, in **Table 1**. Notably, the dataset for testing ANTS was created by selecting the moving and fixed brain images with positional correspondence along the lateromedial axis of the mouse brain at each respective age. All training experiments were performed on an NVIDIA TITAN RTX GPU. For more details on our experimental setup and model architecture, please refer to the Appendix.

**Table 1:** A variety of commonly used deep learning-based segmentation models, SeBRe and U-Net are trained on various combinations of source datasets and tested on unseen target domains. In addition, ANTs affine and non-rigid transformations are applied to compare the performance of the image registration task across domains. The mean DICE coefficient and MSE scores are reported for each model.

| Source(NeuN)/Target | SeBRe |  | U-Net |  | Fix(NeuN)/Mov | ANTs(Aff) |  | ANTs(Aff+NR) |  |
| --- | --- | --- | --- | --- | --- | --- | --- | --- | --- |
|  | DICE | MSE | DICE | MSE |  | DICE | MSE | DICE | MSE |
| P04+P14 / P56 NeuN | 0.89 | 0.02 | 0.83 | 0.04 | P04 / P14 NeuN | 0.61 | 0.06 | 0.69 | 0.05 |
| P04+P56 / P14 NeuN | 0.91 | 0.19 | 0.89 | 0.02 | P04 / P56 NeuN | 0.54 | 0.07 | 0.64 | 0.06 |
| P14+P56 / P04 NeuN | 0.86 | 0.03 | 0.67 | 0.06 | P14 / P56 NeuN | 0.65 | 0.07 | 0.76 | 0.06 |
| P04+P14 / P56 GAD1 | 0.89 | 0.02 | 0.81 | 0.04 | P04 / P14 GAD1 | 0.58 | 0.07 | 0.65 | 0.05 |
| P04+P56 / P14 GAD1 | 0.90 | 0.02 | 0.82 | 0.03 | P04 / P56 GAD1 | 0.48 | 0.08 | 0.60 | 0.06 |
| P14+P56 / P04 GAD1 | 0.84 | 0.03 | 0.69 | 0.06 | P14 / P56 GAD1 | 0.64 | 0.07 | 0.71 | 0.06 |
| P04+P14 / P56 GAD2 | 0.88 | 0.03 | 0.83 | 0.03 | P04 / P14 GAD2 | 0.57 | 0.07 | 0.63 | 0.05 |
| P04+P56 / P14 GAD2 | 0.90 | 0.02 | 0.75 | 0.05 | P04 / P56 GAD2 | 0.49 | 0.08 | 0.57 | 0.06 |
| P14+P56 / P04 GAD2 | 0.84 | 0.03 | 0.60 | 0.07 | P14 / P56 GAD2 | 0.69 | 0.07 | 0.70 | 0.06 |
| P04+P14 / P56 DAPI | 0.88 | 0.03 | 0.79 | 0.04 | P04 / P14 DAPI | 0.52 | 0.07 | 0.63 | 0.060 |
| P04+P56 / P14 DAPI | 0.90 | 0.02 | 0.79 | 0.04 | P04 / P56 DAPI | 0.49 | 0.08 | 0.60 | 0.06 |
| P14+P56 / P04 DAPI | 0.86 | 0.03 | 0.67 | 0.06 | P14 / P56 DAPI | 0.63 | 0.07 | 0.70 | 0.07 |

**Results**: To demonstrate the cross-age domain generalization performance of all deep-learning-based segmentation and registration models, we report the MSE score and DICE score on the whole-cortex segmentation task and cortical registration tasks, in **Table 1**. SeBRe outperforms most of the tasks, followed by U-Net and ANTs. It is noteworthy that models trained on the combination of P4+P56 data perform significantly better on the unseen P14 test dataset, primarily due to the structural variation coverage at the intermediate age by the young and adult brain sections (P4 and P56). Furthermore, domains that cover prominent features of the cortex in the training set, even smaller in data size, still perform highly on unseen target sets e.g. train on DAPI and test on other domains.

#### 3.6.2 Multi-level domain generalization

**Setup**: This is an advanced task, that aims to assess the performance of segmentation and registration models on multi-tier domain variance in the dataset, incorporating cortical structural variance across different mouse ages, as well as data distribution shifts induced by different brain tissue labeling techniques. Segmentation models, Mask-RCNN and 2D U-Net were trained and tested on the dataset splits detailed in **Table 1**, rows 4-12. Both model variants of ANTS (Affine and Affine+Non-rigid) were also put to test on a similarly designed multi-level domain invariance task, detailed in **Table 1**.

**Results**: To demonstrate the multi-level domain generalization performance of all models, we report the MSE and DICE scores on the whole-cortex segmentation and registration tasks, in **Table 1**.

## 4 Discussion and Conclusion

A set of high-resolution cortical atlases is presented for three landmark-developing mouse postnatal ages, P4, P14, and P56, which is based on gene expression-dependent fine-structure annotation. This neuroinformatics resource would enable deep-level anatomical and functional characterization of the murine brain cortex by enabling precise localization of cortical subregions, across developmental time points that remain understudied at such detail due to the prior lack of fine-annotated cortical atlases for young mouse ages. In addition, a novel human-annotated domain generalization benchmark is presented for fine-grained cortical segmentation and cortical atlas registration tasks. This benchmark incorporates a highly enriched dataset, spanning the structural variance of the cortex across critical mouse developmental time points, as well as the domain variance introduced by different genetic markers and stains for nervous tissue, which are commonly encountered in the neuroimaging community. Furthermore, we pose model evaluation tasks at various degrees of complexity, making this a high-utility benchmark for a range of machine-learning problems.

Although the list of genetic markers used for developing atlas validation was composed to be concise, it is not exhaustive of all the cortical subregions defined in the adult mouse reference atlas, and in the case where none of the chosen genetic markers addresses the boundary of a particular cortical subregion, the boundaries from the pre-registered cortex stage are left intact. Additionally, the limited selection of genetic markers could potentially introduce a bias in the learning of the segmentation and registration algorithms. The domain generalization can also be further extended through the addition of other genetic markers, especially sparse labels of brain tissue, to increase the heterogeneity of the domains. Other limitations are presented by the challenges associated with experimental data collection and large-scale manual annotation, which caps the available brain images for each section.

The MortX benchmark has profound implications for neuroscience and AI research. In neuroscience, MortX provides a necessary and much sought-after tool for understanding mouse brain development at different postnatal ages. This granular level of detail can offer vital insights into neurodevelopmental disorders, contributing to improved diagnostic and therapeutic strategies. While MortX fills in an application gap specific to the mouse brain imaging community, it is also constructed to complement and add value to existing domain adaptive segmentation benchmarks in the medical imaging community (Dorent et al. [2023], Memmel et al. [2021]). In the domain of AI, MortX is a significant step forward in addressing the challenge of domain generalization. The benchmark can be used to evaluate the robustness of machine learning models against shifts in data distribution, contributing to the development of more reliable and domain-invariant models. This benchmark also seeks to reduce the need for extensive annotations by modifying networks that are already trained on established reference data (source domain) to function in a new (target) domain with minimal extra annotations. By releasing MortX as open-source, we hope to stimulate advancements in both neuroscience and AI and in the intersection of the two disciplines.

## 5 Methods

### 5.1 Cortical atlas generation

#### 5.1.1 Mapping adult cortical atlas to the developing mouse brain

The adult mouse brain reference atlas from the open-source Allen Brain Map reference atlases (Atlas [2011]) was used as the spatial coordinate framework for cortical subregions. The anatomical template for this atlas comprises 21 brain sections of the left hemisphere of a male mouse brain dissected at postnatal day 56, cut in 25 *µ*m-thick sagittal planes that are spaced 200 *µ*m apart. The brain sections are stained for Nissl substance, which is the ribosomal RNA associated with rough endoplasmic reticulum. Nissl staining highlights cytoarchitectural differences and delineates boundaries between distinct brain tissue types.

The target atlas for the P4 mouse brain was obtained from the open-source Allen Brain Map reference atlases. The anatomical template for this atlas comprises 23 brain sections of the left hemisphere of a male mouse brain at postnatal day 4, cut in 20 *µ*m-thick sagittal planes that are spaced 160 *µ*m apart. The brain tissue sections are labeled with Nissl stain. The target atlas for the P14 mouse brain was obtained also from the open-source Allen Brain Map reference atlases. The anatomical template for this atlas comprises 39 brain sections of the left hemisphere of a male mouse brain at postnatal day 14, cut in 25 *µ*m-thick sagittal planes that are spaced 200 *µ*m apart. The brain tissue sections are labeled with Nissl stain. The sections of the adult mouse brain reference atlas were aligned with their equivalent sections in the developing mouse brain reference atlas at the ages P4 and P14, correspondingly. One-to-one mapping of adult reference brain sections onto positionally-matching developing mouse brain sections is shown in **Supplementary Figure 1**.

An SVG file of a P56 reference brain section was superimposed on the corresponding image of the P4/P14 reference brain section. The outer edge of the P56 reference cortex was aligned with the cortical boundary of the developing mouse reference section beneath it, resulting in the *pre-registered cortex*. This was accomplished through fine-tuning each vector that marked the boundaries of cortical subregions while ensuring the least possible modification to the original shapes and size-ratios of the cortical subregions. The registration process for external boundary alignment is illustrated in **Supplementary Figure 2**.

#### 5.1.2 Cortical gene expression-dependent arealization and layerization

After registration of the P56 mouse cortex to the P4 and P14 cortex, the cortical subregion boundaries were validated using genetic markers that have area-specific and/or layer-specific expression. In-situ hybridized (ISH) brains labeled with the selected markers, were obtained from the open-source Allen Developing Mouse Brain Atlas database. The *pre-registered cortex* was overlaid with each ISH brain section. The ISH reference sections were transformed manually, so as to maximize the alignment between the external boundary of the *pre-registered cortex* and the cortical boundary of the ISH section. This process of coarse registration of each genetically-labelled reference brain image to the *pre-registered cortex* is shown in **Supplementary Figure 3**. Subsequently, the demarcations of the cortical subregions are refined following the expression profile of genetic markers in the matched reference ISH brain image, through careful vector adjustment within the already registered cortex section. The method for precise alignment of cortical subregions is depicted in **Supplementary Figure 4**. In addition to the ISH brain images for the selected neuronal markers, their corresponding gene expression heat maps were also used to aid localization of the structure-specific gene expression **Supplementary Figure 5**.

#### 5.1.3 Genetic markers for boundary alignment

The genetic neuronal markers for the developing brain that were utilized to localize cortical subregion boundaries (cortical arealization and cortical layerization) are described below in further detail.

*P4*

*Arealization:*

**Bhlhe22** (Basic Helix-Loop-Helix Family Member E22) is a transcription factor expressed post-mitotically in glutamatergic neurons of layers 2-5 of the developing cortex. It is expressed at a high caudomedial to low rostrolateral gradient and has been identified as a marker of area-identity between the sensory and motor cortical areas at mouse age P4. Bhlhe22 particularly labels the primary somatosensory, auditory, and visual areas at P4 (Joshi et al. [2008], Woodworth et al. [2012]). The expression of Bhlhe22 is also downregulated at the areal boundary between the neocortex and the cingulate cortex, defining a distinct border at P4 (Gilbert and Ng [2018]).

**Rorb** (RAR-related orphan receptor beta) is a transcription factor with graded rostrocaudal and lateromedial expression in the developing neocortex. At P4, this graded expression is disjuncted at the borders of the primary sensory areas (somatosensory, visual, and auditory cortices)(O’Leary et al. [2007]), allowing delineation of these areal boundaries.

**Cdh6** (Cadherin-6) is a cell adhesion protein that localizes to neuronal synapses. At P4, Cdh-6 is expressed in the primary somatosensory, primary auditory, and motor areas of the mouse cortex, while expression is downregulated in the adjacent cingulate cortex, thus delimiting the boundaries of these cortical regions (Inoue et al. [2008], Hertel and Redies [2010]).

**Lmo4** (LIM Domain Transcription Factor LMO4) and **Cdh8** (Cadherin 8) gene expression data were additionally consulted for the specification of the motor cortex boundaries.

*Layerization:*

**Bhlhe22** is expressed in the cortical layers 2, 3, 4, and 5 with differential expression intensities, co-localizing with other known layer-specific genetic markers. At P4, Bhlhe expression is very sparse in layer 4, which contrasts with the high expression in the flanking layers 2/3 and layer 5 hence defining clear laminar borders. Expression is absent in layer 6, demarcating a conspicuous lower bound for layer 5 (Joshi et al. [2008]).

**Rorb** labeling at P4 is restricted to strong expression in cortical layer 4, with rare co-labeling of layer 5 neurons (Ackman et al. [2014], Roubertoux et al. [2018]), which facilitates the clear isolation of these cortical laminae.

**Cdh6** expression spans layer 2/3 and a belt of deeper layer 4, in the P4 mouse brain(Hertel and Redies [2010]). This marks the laminar borders of layers 2-4.

*P14*

*Arealization:*

**Rorb** expression in cortical layer 4 is downregulated at the borders of the primary sensory areas, highlighting the separation of these areas (Gilbert and Ng [2018]).

*Layerization:*

**Bhlhe22** At P14, the laminar-specific expression of Bhlhe22 across layers 2-5, albeit reduced(Roubertoux et al. [2018]), aids delineation of the cortical laminar borders.

**Bcl6** (B-cell lymphoma 6) is a protein expressed in the long-range projection neurons of layer 5, at P14, distinctly labeling the extent of this cortical lamina. **Rorb** expression at age P14 becomes sharply restricted to layer 4.

Additionally, **Lmo4** and **Cdh8** gene expression data were consulted forthe specification of the motor cortex boundaries (Gilbert and Ng [2018]).

### 5.2 Fine-structure color annotation of developing cortical subregions

After refining the cortical subregion boundaries, guided by genetic marker expression, each cortical subregion was assigned a unique RGB color code. Fine-structure annotated cortical maps for developing brain ages are shown in **Supplementary Figure 6**, for representative lateral brain sections.

### 5.3 Data collection

#### 5.3.1 Animals (genotypes and ages)

Animal experiments were performed following the ARRIVE guidelines. Mice were bred on a C57BL/6 background and kept in standard housing conditions with food and water provided ad libitum. Heterozygous GAD65-cre;GAD67-GFP male mice were bred with heterozygous tdTom female mice (B6.CAG-tdTomato)/J). The GAD65-tdTom;GAD67-GFP offspring were used for immunostaining of the neuronal nuclear marker (NeuN). Three mice from three different litters and both sexes were used per postnatal age (P): 3, 14, and 59.

#### 5.3.2 Tissue extraction

GAD65-Cre;GAD67-GFP animals of both sexes were deeply anesthetized with an intraperitoneal injection of sodium pentobarbital (50 mg/kg), and perfused transcardially with ice-cold phosphate buffer saline (PBS), pH 7.4 to rinse blood, followed by fixative containing 4% paraformaldehyde in 0.15 M Na-phosphate buffer, pH 7.4. Perfusion was done manually for P3 pups using a Butterfly Infusion (27g) connected to a 10 ml syringe and for P14 and P59 animals using a perfusion pump at a constant flow rate of 8-12 ml/min. Brains were immediately dissected, cut sagittal through the midline, and postfixed in ice-cold 4% paraformaldehyde (dissolved in 0.15 M Na-phosphate buffer, pH 7.4) 72 hours for P3, 12 hours for P14 and 3 hours for P59. Brains were then rinsed twice with ice-cold PBS, transferred to 30% sucrose in PBS for cryopreservation, and stored for 3 days at 4°C. Brains were frozen with dry ice and kept at −80°C until further use.

Sagittal sections from one hemisphere were cut into serial sections at 60 *µ*m for P3, and 40-*µ*m thickness for P14 and P59 using a sliding blade freezing microtome (HM400; Microm). Two serial sections were collected for brains at P3 (120 *µ*m serial sampling), five serial sections (200 *µ*m serial sampling) for P14, and 6 serial sections (240 *µ*m serial sampling) for P59. Sections were stored at −20°C in an antifreeze solution until use.

#### 5.3.3 Immunofluorescence staining

Triple immunofluorescence staining was used to analyze multiple neuronal markers (Gad65, Gad67 and NeuN) within the same section. Free-floating sections from GAD65-Cre;GAD67-GFP were washed three times for 10 min each in Tris-Triton buffer, pH 7.4 before being incubated overnight in a dark box at 4°C under continuous agitation with the primary monoclonal mouse nuclear neuronal antibody (NeuN) (MAB377, Chemicon) diluted 1:[1000] in a solution containing 2% Triton X-100 and 2% NGS in Tris-Triton buffer, pH 7.4. The next day, the sections were rinsed three times for 10 min with Tris-Triton buffer, pH 7.4 followed by incubation with secondary antibodies against the mouse coupled to Alexa Fluor 647 in a solution containing 2% NGS in Tris-Triton buffer, pH 7.4 at RT for 30 min in the dark. After another washing step of three times 10 min with Tris-Triton buffer, pH 7.4, sections were mounted on gelatin-coated glass slides, cover-slipped with Dako fluorescence mounting medium (Dako) and stored at 4°C.

#### 5.3.4 Fluorescence imaging

Immunolabelled serial sections were imaged with a whole slide-scanning system, Zeiss Axio Scan.Z1, using an air 20X (NA 0.8) objective and multi-channels for green (excitation 470, beam splitter FT495, emission BP 525/50), red (excitation 555, beam splitter FT570, emission BP 605/70) and far red (excitation 630, beam splitter FT660, emission BP 690/50) fluorescence. The equipment operates with the ZEN Image Analysis module.

After a preview scan of the slide, the tissue detection (TD) area was selected leaving an air gap around the slide so that the scan can detect the edges of the tissue section. Image acquisition was done using automated threshold and focusing settings. Focus spacing was N=3. Images were taken with a Hamamatsu Orca Flash 4.0 monochrome camera 16bit (fluorescence camera), [2048] x [2048] pixels (4 MP), pixel size 6.5 µm.

#### 5.3.5 Allen brain repository data

ISH brains, labeled with GAD1 and Nissl (mouse ages P4 and P14) and co-labeled with CamK2/GAD1 (mouse age P56) were acquired from the open-sourced Allen Developing Mouse Brain repository (Gilbert and Ng [2018]).

### 5.4 Multimodal data distributional shifts

In order to quantitatively demonstrate the domain variation captured by our dataset, we created low dimensional UMAP embeddings of all brain section images labeled with distinct genetic markers and nuclear stains, within a single mouse age group, and project these embeddings to a shared latent space. UMAP reduces each brain section image to a 2-dimensional(2D) representation. These 2D-embeddings are presented separately for each mouse age (P4, P14, and P56), as well as for the complete data dataset, in **Supplementary Figure 26**. Low dimensional embeddings highlight a distributional shift induced by each unique brain marker, within and across mouse ages. Sections belonging to the same brain tend to cluster together, showing within-domain similarity.

In addition, we qualitatively present the variability in brain image attributes, such as luminance, contrast and structure, across different mouse brain ages and brain tissue labels, in **Supplementary Figures 10 - 24**. This emphasizes the richness of our dataset in terms of domain heterogeneity and, thus, its utility as a domain generalization benchmark for the neuroimaging community.

The brain images available for each domain are enumerated in **Supplementary Table 2**.

### 5.5 Models for cortex segmentation and registration

**SeBRe**: We used the SeBRe model (Iqbal et al. [2019]), an extension of the Mask R-CNN, known for its efficiency in instance segmentation tasks. We implemented the model using the PyTorch framework. The configuration of our Mask R-CNN involved a ResNet-101 backbone for feature extraction, and the Region Proposal Network (RPN) anchor scales were set at [32, 64, 128, 256, 512], with anchor aspect ratios of [0.5, 1, 2]. RoIAlign was utilized for maintaining spatial precision while extracting features from the proposed regions. The training procedure followed used stochastic gradient descent (SGD) as the optimization method with an initial learning rate of 0.[0002]5. The model was trained for [1000] epochs with a batch size of 2, using a weight decay of 0.[0001] and a momentum of 0.9 for regularization.

**U-Net**: We utilized the U-Net model(Ronneberger et al. [2015]), widely recognized for its remarkable efficiency in biomedical image segmentation tasks, for our study. The model was implemented in a Python 3.8 environment using the TensorFlow framework (version 2.6.0). Our U-Net configuration comprised an encoding (contraction) path and a symmetric decoding (expansion) path, creating the namesake “U” shape. The encoding path consisted of repeated applications of two 3×3 convolutions followed by a ReLU activation function and a 2×2 max pooling operation. In the decoding path, a similar repeated structure of operations was used, including 2×2 up-convolution, concatenation with the corresponding feature map from the encoding path, followed by two 3×3 convolutions and a ReLU activation function. The encoder and decoder modules consist of 5 conv blocks. The model was trained using the Adam optimizer with a learning rate of 0.[0001] for 100 epochs, with a batch size of 1. Model regularization was performed via early stopping based on the validation loss.

**ANTs**: We employed the Advanced Normalization Tools (ANTs)(Avants et al. [2011]), a widely recognized computational framework known for its high performance in the registration and normalization of medical imaging data. We utilized the symmetric normalization (SyN) algorithm, a core part of the ANTs suite, which enables high-dimensional nonlinear registration via diffeomorphic transformations. In the process, we configured the algorithm to use a cross-correlation similarity metric and a multi-resolution scheme. The gradient step was set to 0.25 with ‘SyNOnly’ as the type of transform and registration iterations of (10,10,10,10) in affine and (256,256,256) in non-rigid, respectively. The flow and total sigmas were set to 9 and 0.2 with the nearest neighbor as the default interpolator.

The performance of all models was evaluated using using mean squared error (MSE) and DICE similarity coefficient. All training and testing were conducted on hardware equipped with an NVIDIA TITAN RTX GPU.

### 5.6 Benchmarking with domain adaptive methods

Based on the superior segmentation performance of Mask RCNN (see main paper) on domain generalization tasks, we elect Mask-RCNN to demonstrate the utility of our dataset on domain generalization tasks, after the brain section images have been preprocessed with commonly used domain normalization techniques: local normalization (LN)(Sage and Unser [2001]), local contrast normalization (LCN)(Jarrett et al. [2009]), local response normalization (LRN)(Krizhevsky et al. [2012]), and z-score normalization. These adaptive techniques are typically applied during image processing, in order to rectify a distribution shift between the source and target domains and to increase domain homogeneity in the data. This step allows downstream image analysis and image-manipulation tasks to be performed on the pre-treated dataset in a domain-invariant manner. Qualitative examples of domain adaptation by the above-listed techniques, across different brain markers, are shown in **Supplementary Figure 25**. It can be observed that these techniques tend to minimize the distributional shift across the different imaging domains and highlight the generalized features of brain regions (including the cortex). LN is observed to filter out the low-frequency pixels whereas LRN focuses on enhancing constrastive features.

### 5.7 Domain generalization across cortical developmental time-points

**Setup.** In this task, whole-cortex segmentation is performed on brain images labeled with the panneuronal genetic marker (NeuN) and pre-processed using the above-mentioned domain adaptive techniques. We evaluate the cross-age structural invariance of Mask-RCNN in combination with each domain adaptive technique. All training (source domain) and testing (target domain) dataset splits are detailed under rows 1-3 in **Supplementary Table 1**.

**Results.** We evaluate the segmentation performance of Mask-RCNN using MSE and Dice coefficient. Mask-RCNN in combination with each domain adaptive technique, shows comparable high segmentation performance on all three source/target domain sets testing structural invariance, as seen across rows 1-3 of **Supplementary Table 1**. Notably, applying LRN to the source and target domains increases the segmentation performance of Mask-RCNN across all cross-age source/target sets, highlighting a significant domain shift between these source and target datasets, which is bridged by applying domain adaptation methods. Similar performance enhancements are observed in the case of LN and LCN applications, for domain generalization from P14+P5 brains to P4 brains. This emphasizes that generalizing from younger mouse brains (source domain) to adult mouse brains (target domain) is a high-complexity task.

### 5.8 Multi-level domain generalization

**Setup.** This task aims to assess the segmentation performance of Mask-RCNN on multi-tier domain variance in the dataset, including cortical structural variance across different mouse ages, as well as data heterogeneity introduced by diverse brain tissue labeling techniques and genetic markers. Mask-RCNN was trained and tested on the dataset splits detailed in **Supplementary Table 1**, rows 4-12. Each experiment was performed after athe pplication of domain adaptation techniques to both source/target sets.

**Results.** To demonstrate the multi-level domain generalization performance of all models, we report the MSE and DICE scores on the whole-cortex segmentation task, in **Supplementary Table 1**. Applying domain adaptive pre-processing, LRN and LCN in particular, results in improved segmentation performance over the Mask-RCNN baseline (see main paper, as well as **Supplementary Figure 7** - **Supplementary Figure 9**) across selective source/target domain sets, as highlighted in **Supplementary Table 1**. This underscores a significant domain shift between the source and target datasets, marking these as high-complexity domain generalization tasks. Additionally, the qualitative performance of Mask-RCNN is illustrated on sample brain images, in **Supplementary Figure 7** - **Supplementary Figure 9**.

It is important to note that the genetic marker NeuN is chosen for these experiments as a representative source domain in the above domain generalization tasks. Essentially, all benchmarking tasks detailed in the main and supplementary text can be performed using brain images labelled with any genetic label provided in the MortX dataset.

### 5.9 Within-domain segmentation performance

In order to provide reference scores for a within-domain task, we designed an experiment where Mask-RCNN was trained, with cross-validation (k=3), on two P4 animal brains fluorescently labeled with the inhibitory neuron marker, GAD1, and tested on a third P4 animal brain also fluorescently labeled with GAD1. We report the DICE scores for each experiment in **Supplementary Table 3**.

---

**Supplementary Table 1:** Our best performing deep learning-based segmentation model, SeBRe, in combination with commonly used domain adaptation techniques, LN is trained on various combinations of source datasets and tested on unseen target domains. The mean DICE coefficient and MSE scores are reported for each model. All experiments where applying domain adaptation techniques enhances (based on DICE) domain generalization performance over the baseline SeBRe (see main paper) are marked in boldface.

| Source(NeuN)/Target | SeBRe(LN) |  | SeBRe(LCN) |  | SeBRe(LRN) |  | SeBRe(ZSCORE) |  |
| --- | --- | --- | --- | --- | --- | --- | --- | --- |
|  | DICE | MSE | DICE | MSE | DICE | MSE | DICE | MSE |
| P4+P14 / P56 NeuN | 0.868 | 0.027 | 0.888 | 0.022 | **0.902** | 0.020 | 0.867 | 0.026 |
| P4+P56 / P14 NeuN | 0.899 | 0.024 | 0.908 | 0.020 | **0.917** | 0.018 | 0.899 | 0.023 |
| P14+P56 / P4 NeuN | **0.868** | 0.023 | **0.871** | 0.024 | **0.880** | 0.023 | 0.862 | 0.028 |
| P4+P14 / P56 GAD1 | 0.876 | 0.026 | 0.876 | 0.026 | 0.875 | 0.026 | 0.859 | 0.028 |
| P4+P56 / P14 GAD1 | **0.909** | 0.022 | 0.895 | 0.024 | **0.908** | 0.021 | 0.893 | 0.025 |
| P14+P56 / P4 GAD1 | 0.842 | 0.029 | **0.857** | 0.026 | **0.858** | 0.026 | 0.842 | 0.032 |
| P4+P14 / P56 GAD2 | **0.892** | 0.024 | 0.877 | 0.025 | **0.894** | 0.024 | 0.845 | 0.030 |
| P4+P56 / P14 GAD2 | **0.910** | 0.022 | **0.903** | 0.022 | **0.909** | 0.022 | 0.897 | 0.024 |
| P14+P56 / P4 GAD2 | 0.842 | 0.029 | **0.859** | 0.027 | **0.863** | 0.025 | **0.852** | 0.029 |
| P4+P14 / P56 DAPI | 0.857 | 0.029 | 0.875 | 0.026 | 0.880 | 0.026 | 0.852 | 0.030 |
| P4+P56 / P14 DAPI | 0.899 | 0.024 | **0.920** | 0.018 | **0.913** | 0.020 | 0.887 | 0.026 |
| P14+P56 / P4 DAPI | 0.849 | 0.024 | **0.872** | 0.025 | **0.870** | 0.024 | 0.863 | 0.029 |

**Supplementary Table 2:**
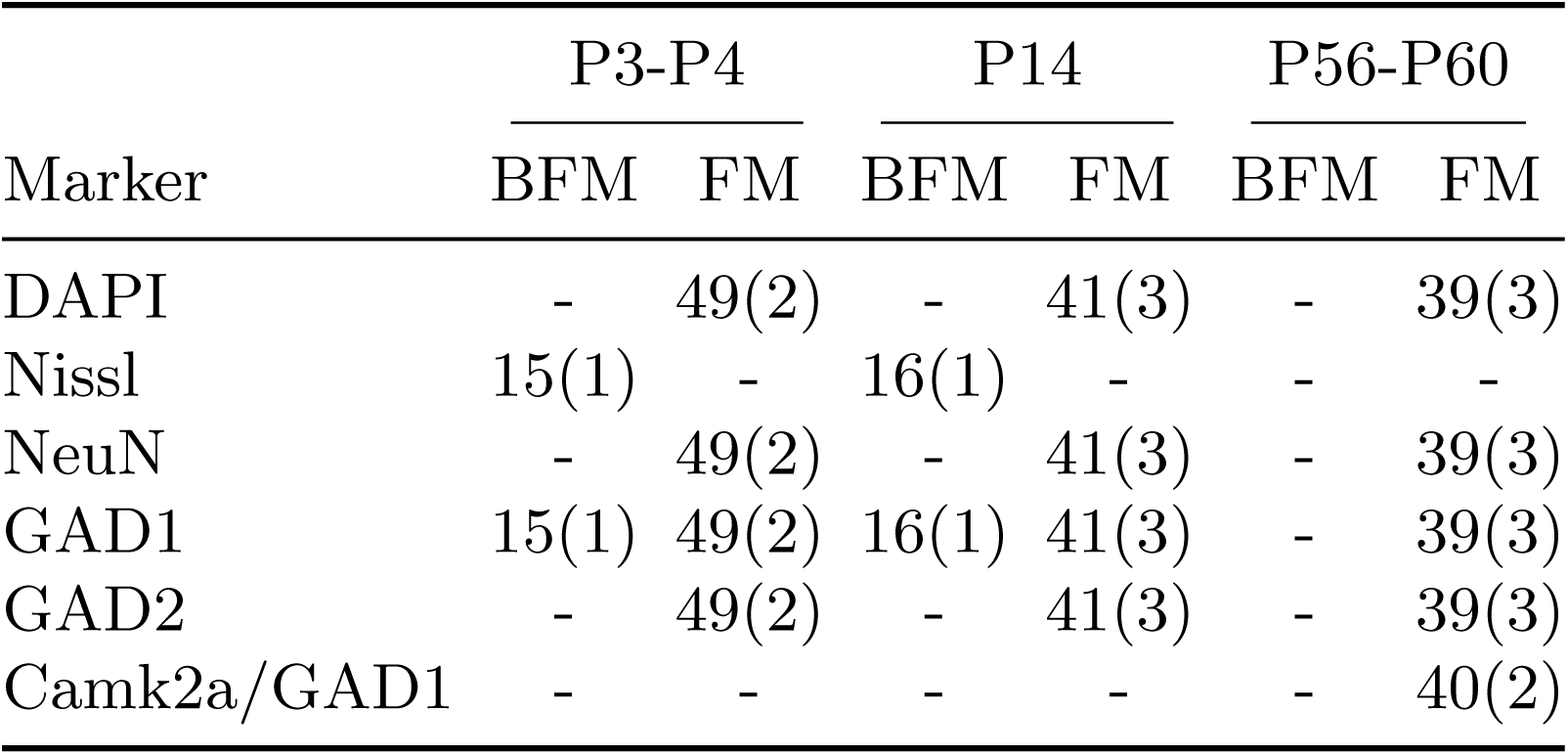
Dataset domain distribution is quantitatively described. The number of brain sections and animals (in parentheses) are enumerated for each genetic marker, imaging modality and mouse age.

**Supplementary Table 3:** SeBRe segmentation performance on a within-domain task. All training and testing images belonged to P4 mice, labelled with a unique fluorescent inhibitory neuron marker, GAD1. The DICE scores for each experiment are report, in addition to the mean and standard error.

| Experiment | DICE |
| --- | --- |
| Trained on animals 1 and 2 — Tested on animal 3 | 0.89 |
| Trained on animals 1 and 3 — Tested on animal 2 | 0.90 |
| Trained on animals 2 and 3 — Tested on animal 1 | 0.95 |
| Mean | $0.92 \pm (0.025)$ |

**Supplementary Figure 1:**
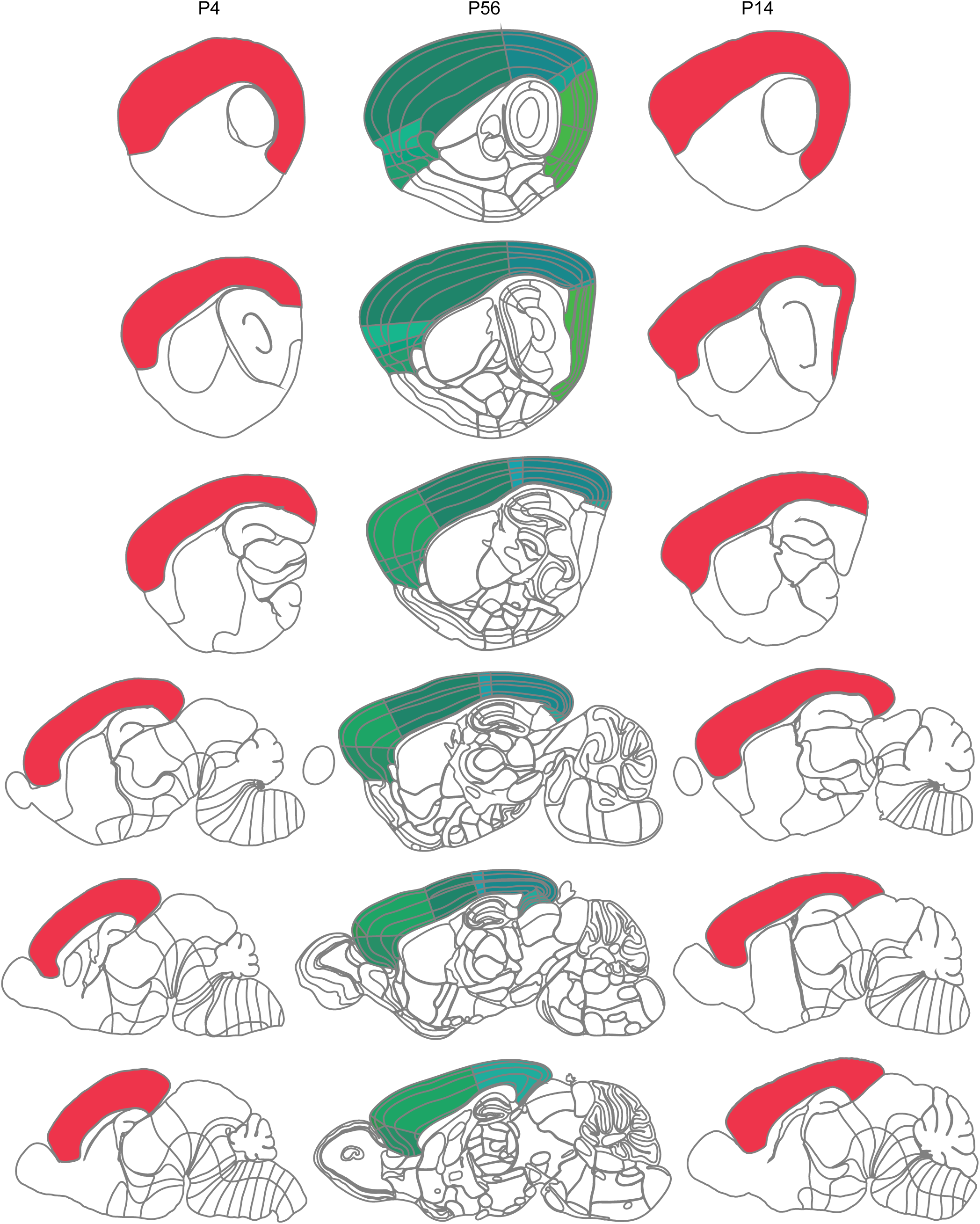
Mapping adult reference brain sections onto positionally-matching developing mouse brain sections. The first and third columns show reference brain sections drawn from different lateral and medial planes of the Allen Brain atlases for the P4 and P14 mouse, respectively. The second column shows the selected reference sections at the corresponding lateromedial planes of the Allen P56 mouse brain atlas. The selection method maximes similarity between the anatomical boundary of the isocortex (filled), as well as the two-dimensional appearance of other brain regions (outlined) in that reference plane.

**Supplementary Figure 2:**
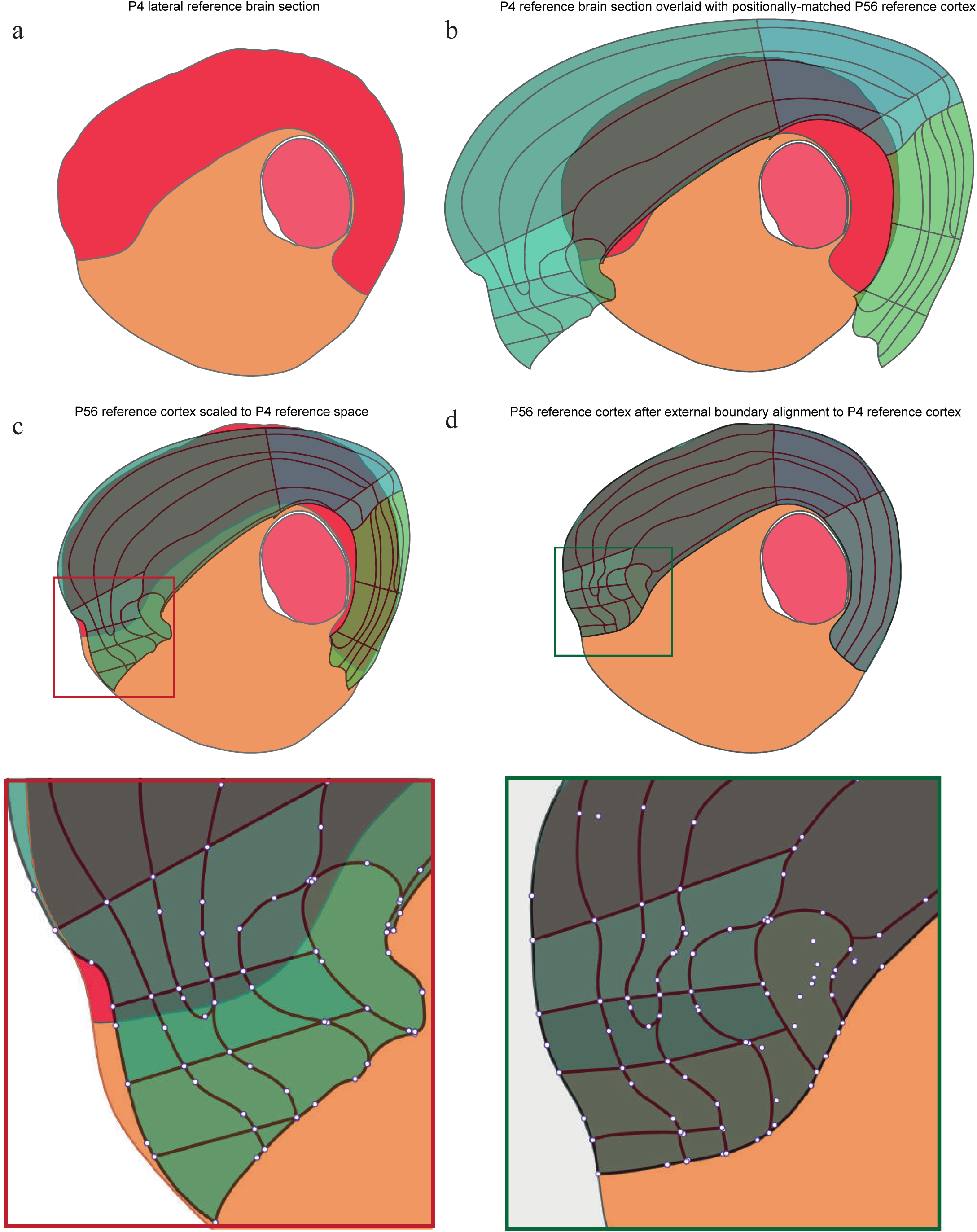
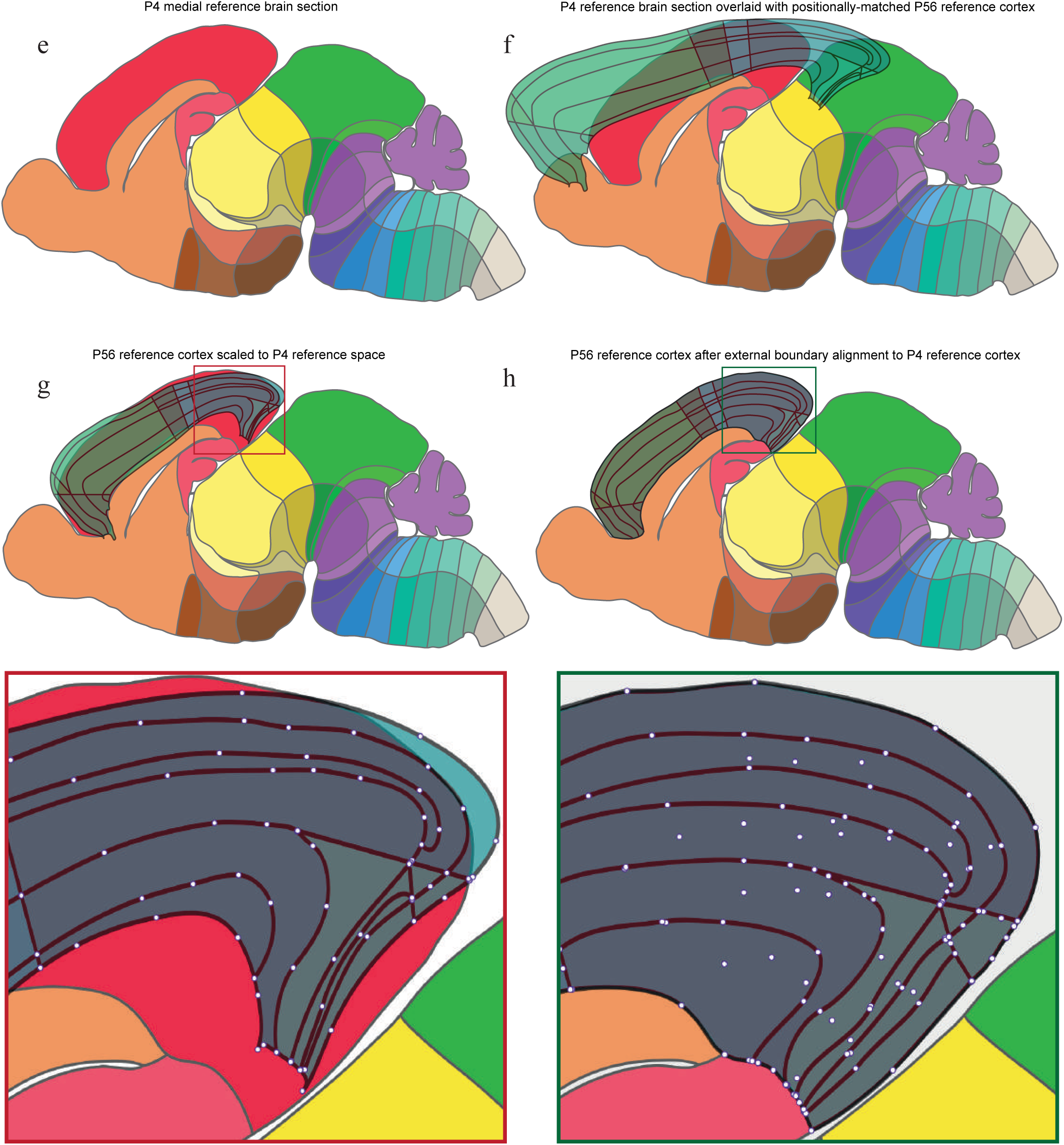
External boundary registration of the adult mouse cortex to the developing cortex reference space. a,e Example lateral (a) and medial (e) reference brain sections of the developing P4 mouse cortex are shown. b,f The lateral (b) and medial (f) reference brain atlas sections of the P4 mouse brain are overlaid with the corresponding positionally-matched P56 lateral and medial reference cortical atlas sections, respectively. c,g The overlaid P56 cortical atlas is scaled to the reference space of the P4 mouse atlas. The inset (red) shows a detailed view of the vector positions (white circles), which define the laminar and areal borders, of the overlaid adult cortex after scaling. d,h The overlaid P56 mouse cortical atlas is shown after complete external boundary alignment to the cortex of the underlying developing mouse brain sections. The inset (green) shows a detailed view of the vector positions of the overlaid adult cortex after complete external boundary alignment, through manual application of a form of non-rigid transformation.

**Supplementary Figure 3:**
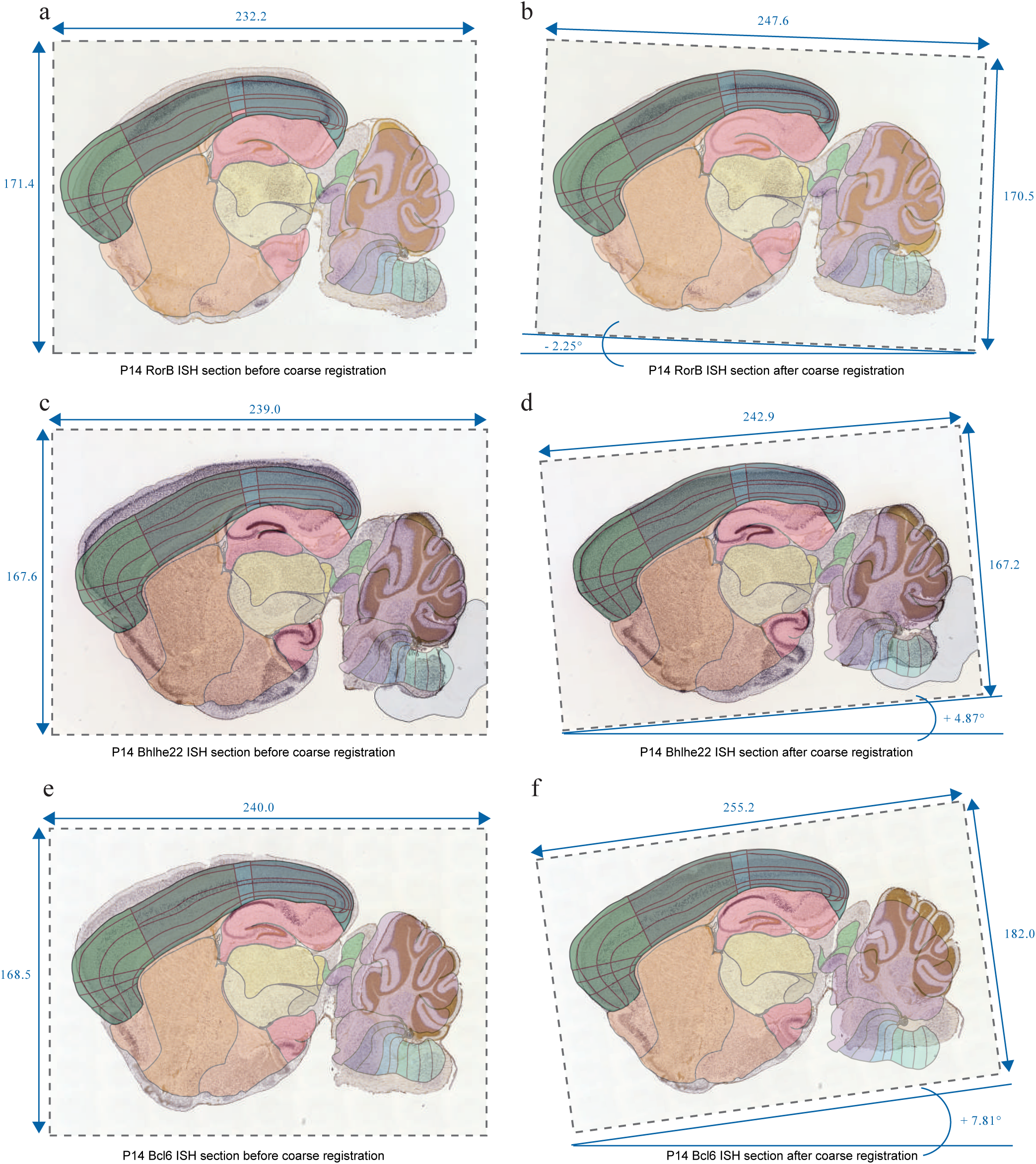
Coarse registration of laminar-specific and area-specific genetic marker-labelled brain images to the pre-registered cortex section. (a-f) ISH brain sections labelled for RorB, Bhlhe22 and Cdh6 mRNA are independently registered to an example P14 mouse pre-registered cortex section. a,c,e The pre-registered cortex section is shown overlaid with an ISH brain section labelled for the neuronal markers RorB (a), Bhlhe (c) and Bcl6 (e), respectively, before registration. The original dimensions of the unregistered ISH section are marked (arbitrary units). b,d,f The pre-registered cortex section is shown overlaid with an ISH brain section labelled for the neuronal markers RorB (b), Bhlhe (d) and Bcl6 (f), respectively, after manual application of a form of non-rigid transformation. The dimensions of the registered ISH brain sections are shown, as well as the applied rotation relative to the original brain section image.

**Supplementary Figure 4:**
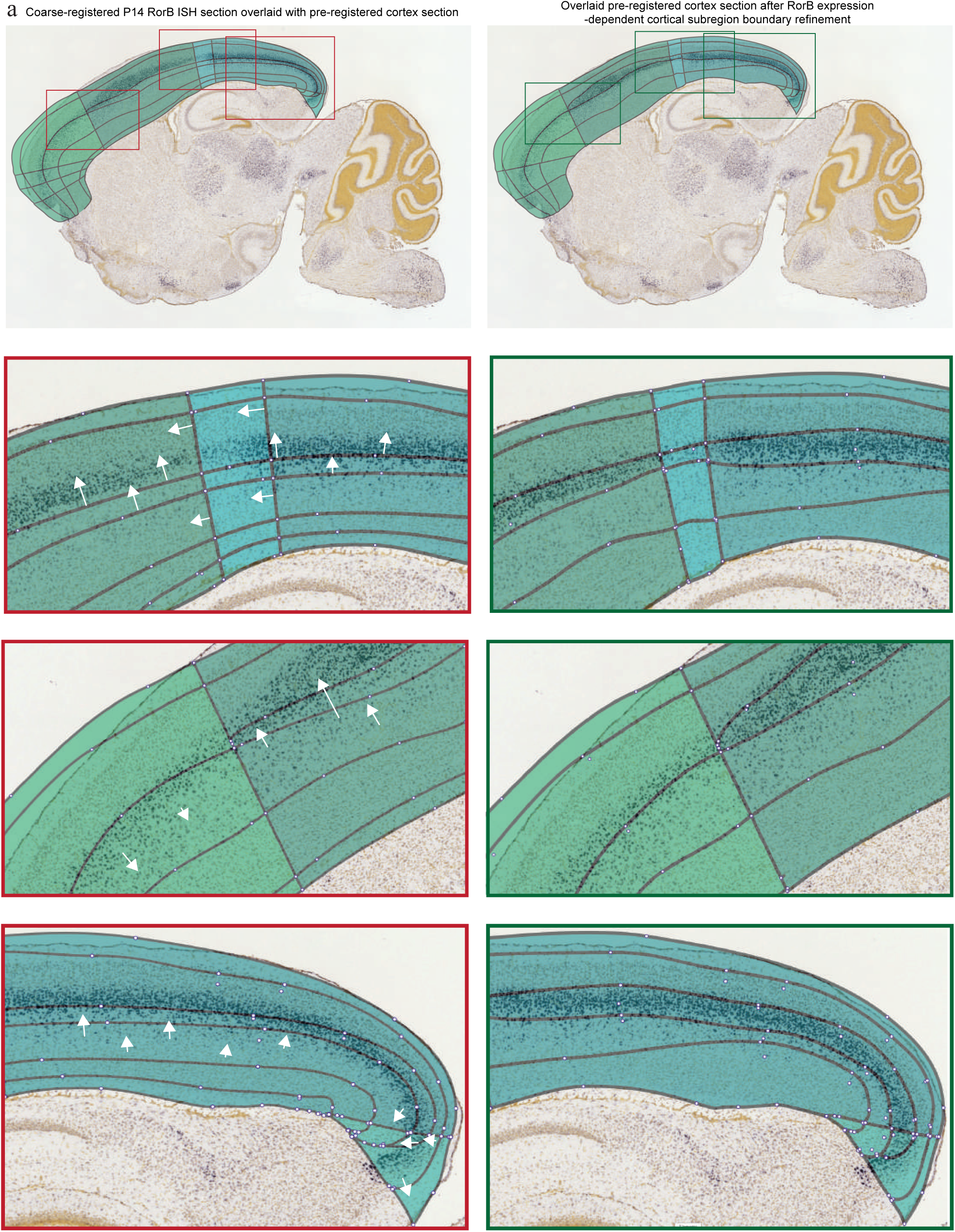
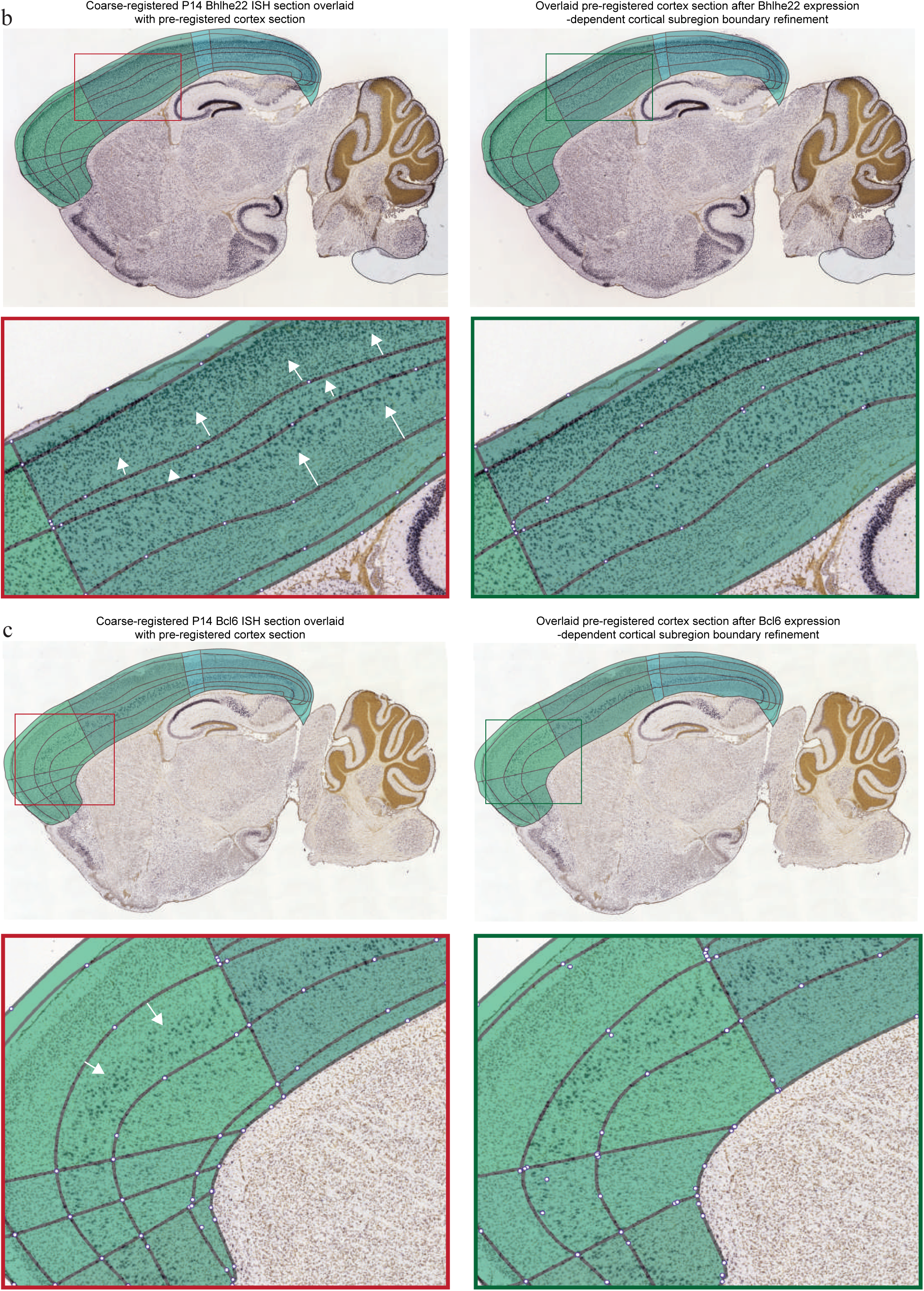
Gene expression-dependent fine registration of internal laminar and areal boundaries of cortical subregions in the pre-registered cortex section. (a-c) Fine structure annotation of cortical subregions in an example P14 mouse pre-registered cortex section, guided by the gene expression intensity of layer-specific and region-specific proteins: RorB, Bhlhe22 and Bcl6. a, First row: the pre-registered cortex section is overlaid onto a coarse-registered ISH brain section labelled for RorB before (left panel) and after (right panel) gene expression-dependent refinement of cortical subregion boundaries. Second row: the left inset (red) shows a detailed view of RorB-guided vector adjustment, illustrated by the white arrows; arrowhead indicates the direction and arrow length indicates the extent of gene expression-guided laminar border modification for layer 4, and the areal border specification between the somatosensory and visual cortical areas. The right inset (green) shows the updated laminar and areal borders. Third row: the left inset (red) shows a detailed view of RorB-guided vector adjustment, white arrows indicate the gene expression-guided laminar border specification of layer 4. The downregulation of RorB expression marks the areal border between the motor and somatosensory cortical areas. The right inset (green) shows the updated laminar borders. Fourth row: the left inset (red) shows RorB-guided vector adjustment, white arrows indicating the gene expression-guided laminar border specification of layer 4. The right inset (green) shows the updated laminar borders. b, First row: the pre-registered cortex section is overlaid onto a coarse-registered ISH brain section labelled for Bhlhe22 mRNA before (left panel) and after (right panel) gene expression-dependent modification of cortical subregion boundaries. Second row: the left inset (red) shows Bhlhe22-guided vector adjustment, white arrows indicating the gene expression-dependent laminar border specification of layer 2/3, layer 4 and layer 5 (Notice distinct gene expression bands in layer 2/3 and layer 5, as well as significant downregulation of expression in the flanked layer 4). The right inset (green) shows the updated laminar borders. c, First row: the pre-registered cortex section is overlaid onto a coarse-registered ISH brain section labelled for Bcl6 mRNA before (left panel) and after (right panel) gene expression-dependent modification of cortical subregion boundaries. Second row: the left inset (red) shows Bcl6-guided vector adjustment, white arrows indicating the gene expression-dependent laminar border specification of layer 5. The right inset (green) shows the updated laminar borders.

**Supplementary Figure 5:**
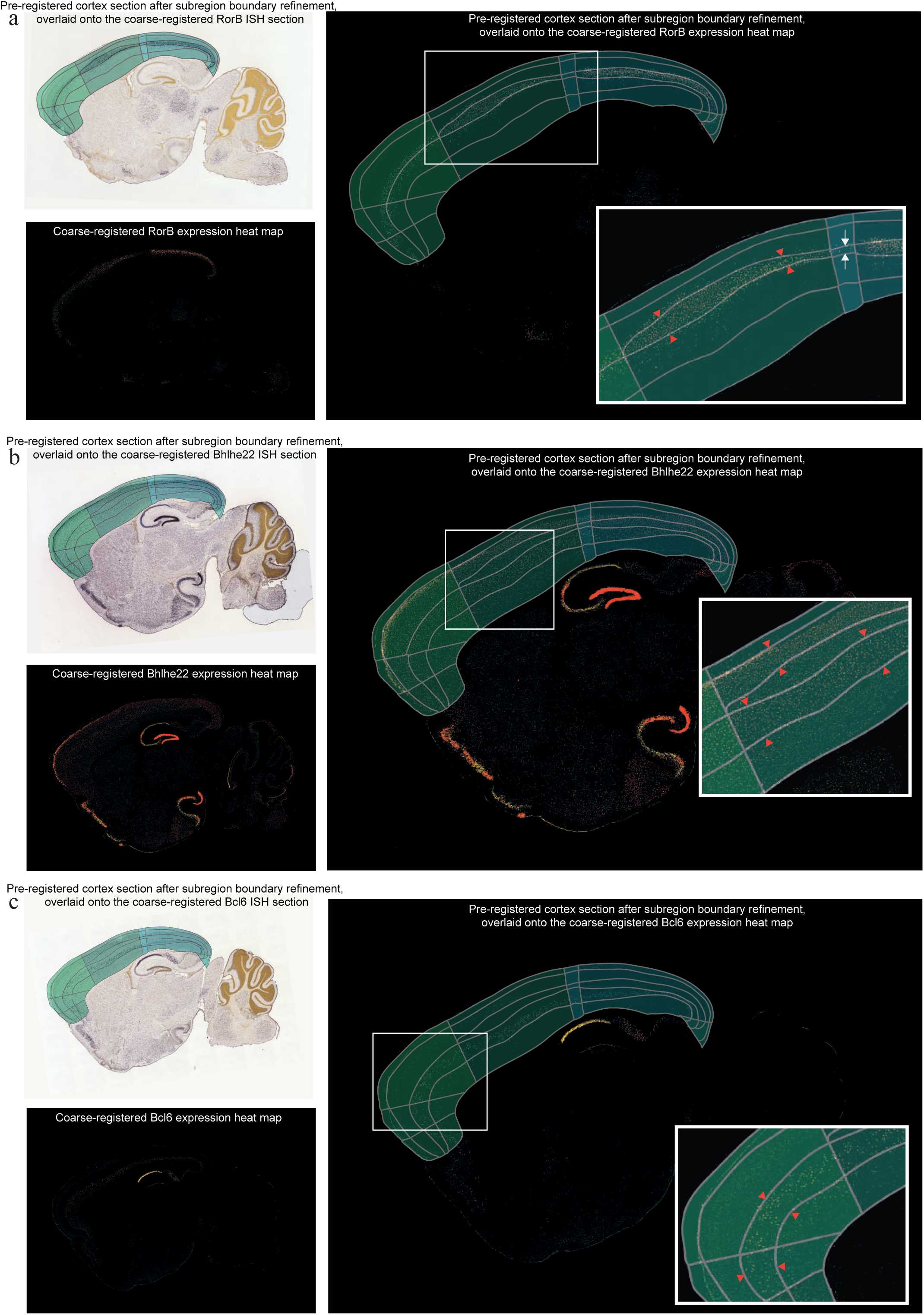
Expression map-aided localization of structure-specific gene expression. a, Fine-scale registration of cortical subregions was validated with gene expression intensity heat maps corresponding to the ISH brain images for structure specifically-enriched RorB, Bhlhe22 and Bcl6 markers. a, Top-left panel shows an example P14 pre-registered cortex section overlaid onto a coarse-registered ISH brain section labelled for RorB, after RorB-guided laminar and areal boundary specification. Bottom left panel shows the gene expression heat map for the coarse-registered RorB ISH data above. Right panel shows the pre-registered cortex section overlaid onto the coarse-registered RorB expression intensity image, after RorB-guided layer 4 delineation, which is detailed in the inset (Orange arrowheads highlight the alignment of gene expression intensity boundaries with the post fine-registration cortical subregion boundaries; white arrows indicate the downregulation in RorB gene expression at the boundary between the somatosensory and visual cortical areas). b, Top-left panel shows the pre-registered cortex section in (a) overlaid onto a coarse-registered ISH brain section labelled for Bhlhe22, after Bhlhe22-guided laminar boundary specification. Bottom left panel shows the gene expression heat map for the coarse-registered Bhlhe22 ISH data in the above panel. Right panel shows the pre-registered cortex section overlaid onto the coarse-registered Bhlhe22 expression intensity image, after Bhlhe22-guided layer 2/3 and layer 5 specification. The inset shows a detailed view of expression intensity-aided localization of laminar borders. c, Top-left panel shows the pre-registered cortex section in (a) overlaid onto a coarse-registered ISH brain section labelled for Bcl6, after Bcl6-guided laminar boundary specification. Bottom left panel shows the gene expression heat map for the coarse-registered Bcl6 ISH data in the above panel. Right panel shows the pre-registered cortex section overlaid onto the coarse-registered Bcl6 expression intensity image, after Bcl6-aided layer layer 5 specification, highlighted in the inset.

**Supplementary Figure 6:**
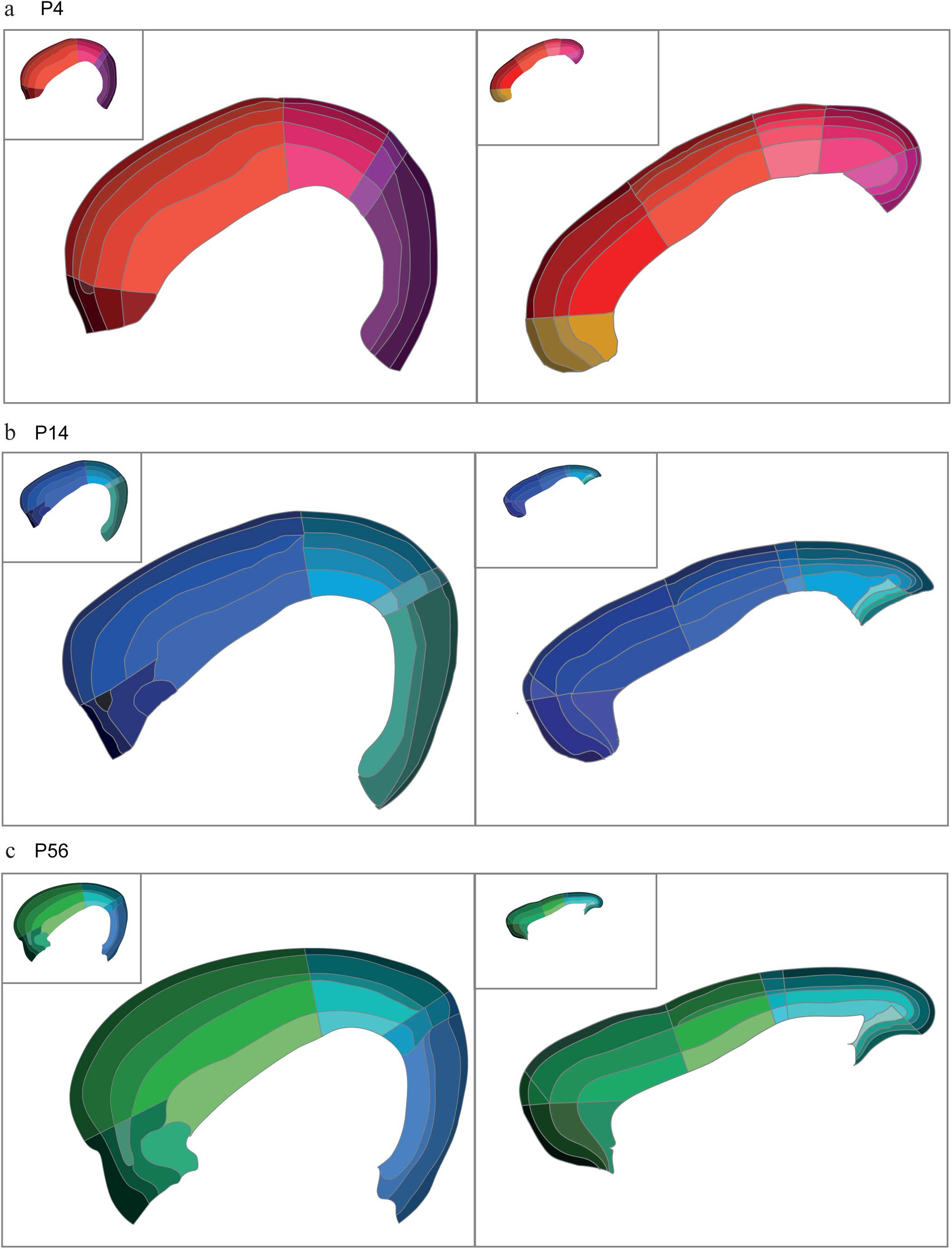
Detailed view of fine-structure annotation of developing mouse cortex. (a-c) Enlarged view of the P4 (a), P14 (b) and P56 (c) cortical reference atlas, drawn from representative lateral (left panel) and medial (right) planes.

**Supplementary Figure 7:**
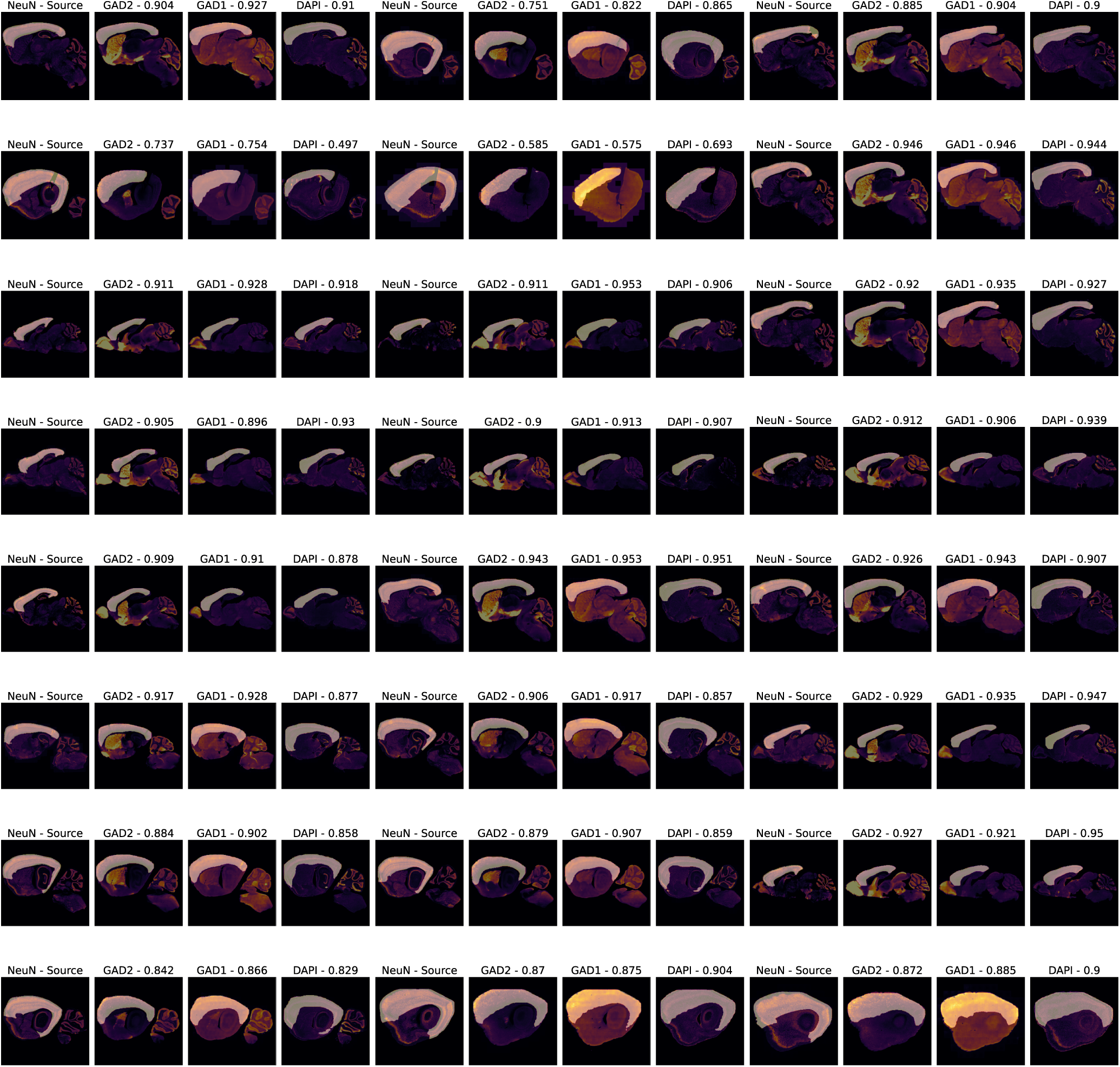
Qualitative and quantitative results on whole-cortex segmentation using Mask-RCNN. The source domain comprises of P4 and P14 brains labelled with NeuN (columns 1, 5 and 9). The target domains consists of P56 brains labelled with GAD1 (columns 3, 7 and 11), GAD2 (columns 2, 6 and 10) or DAPI (columns 4, 8, and 12). Each brain section is overlaid with the predicted whole-cortex segmentation mask. The DICE scores are reported for each input sample.

**Supplementary Figure 8:**
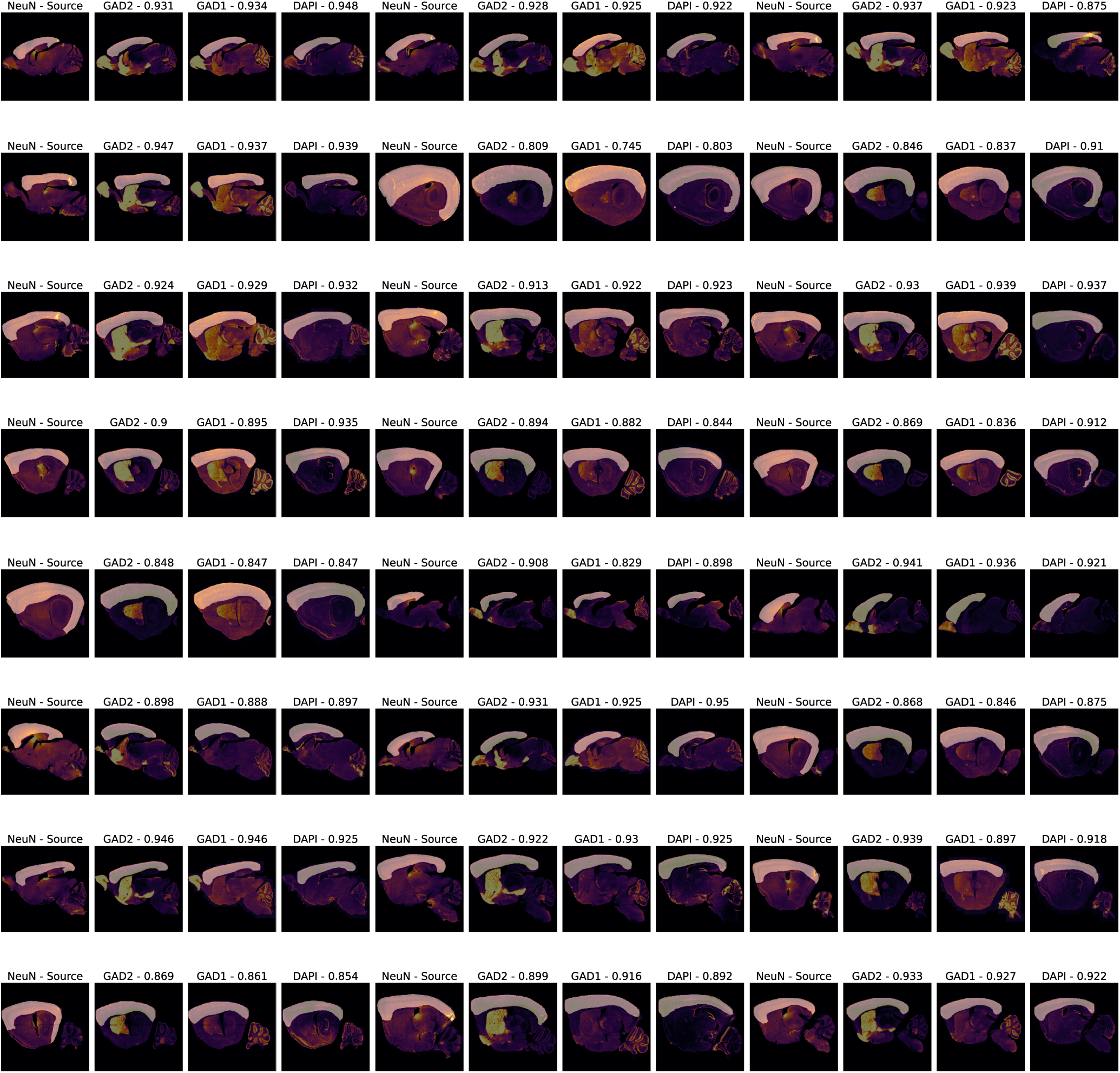
Qualitative and quantitative results on whole-cortex segmentation using Mask-RCNN. The source domain comprises of P4 and P56 brains labelled with NeuN (columns 1, 5 and 9). The target domains consists of P14 brains labelled with GAD1 (columns 3, 7 and 11), GAD2 (columns 2, 6 and 10) or DAPI (columns 4, 8, and 12). Each brain section is overlaid with the predicted whole-cortex segmentation mask. The DICE scores are reported for each input sample.

**Supplementary Figure 9:**
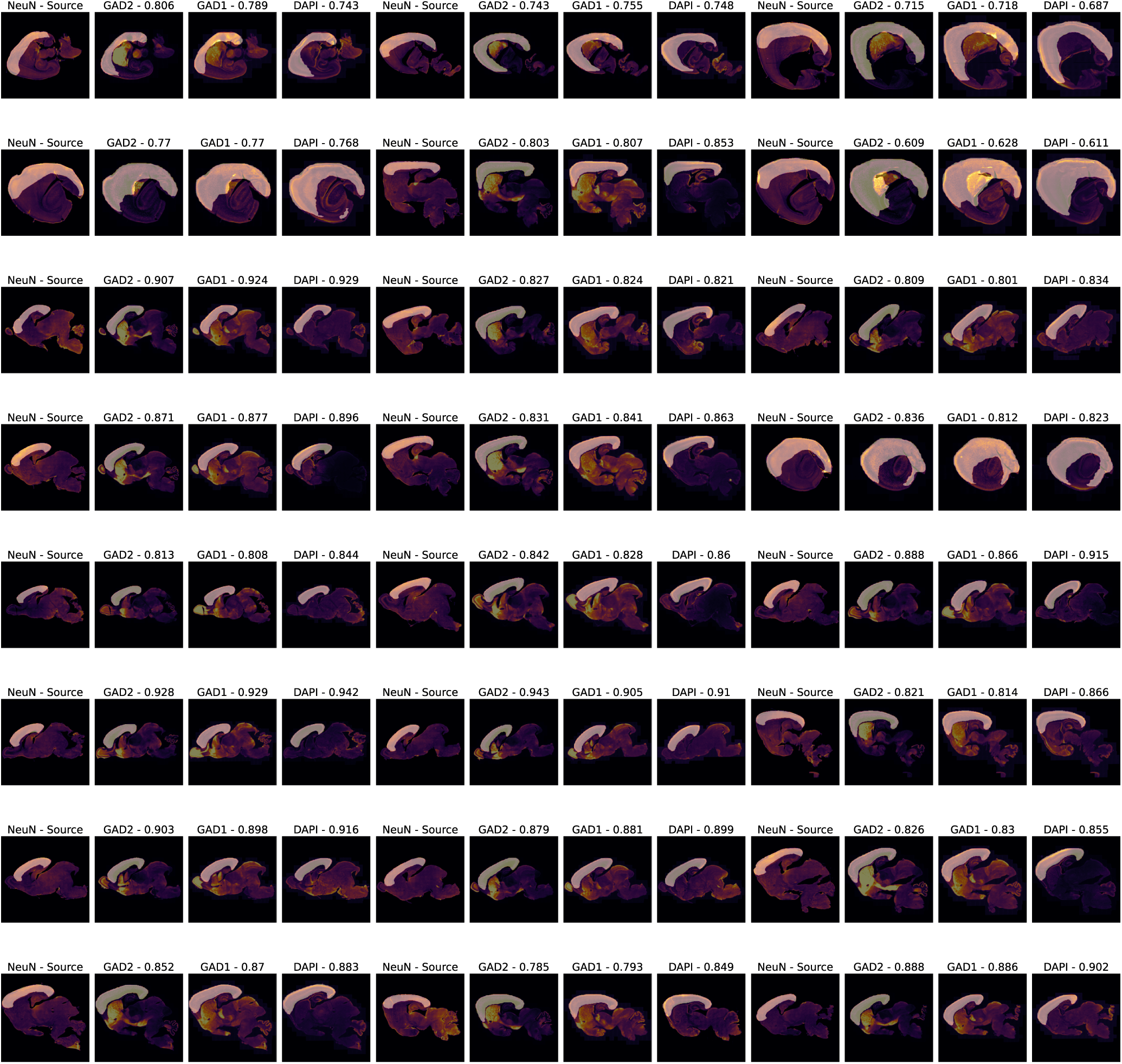
Qualitative and quantitative results on whole-cortex segmentation using Mask-RCNN. The source domain comprises of P14 and P56 brains labelled with NeuN (columns 1, 5 and 9). The target domains consists of P4 brains labelled with GAD1 (columns 3, 7 and 11), GAD2 (columns 2, 6 and 10) or DAPI (columns 4, 8, and 12). Each brain section is overlaid with the predicted whole-cortex segmentation mask. The DICE scores are reported for each input sample.

**Supplementary Figure 10:**
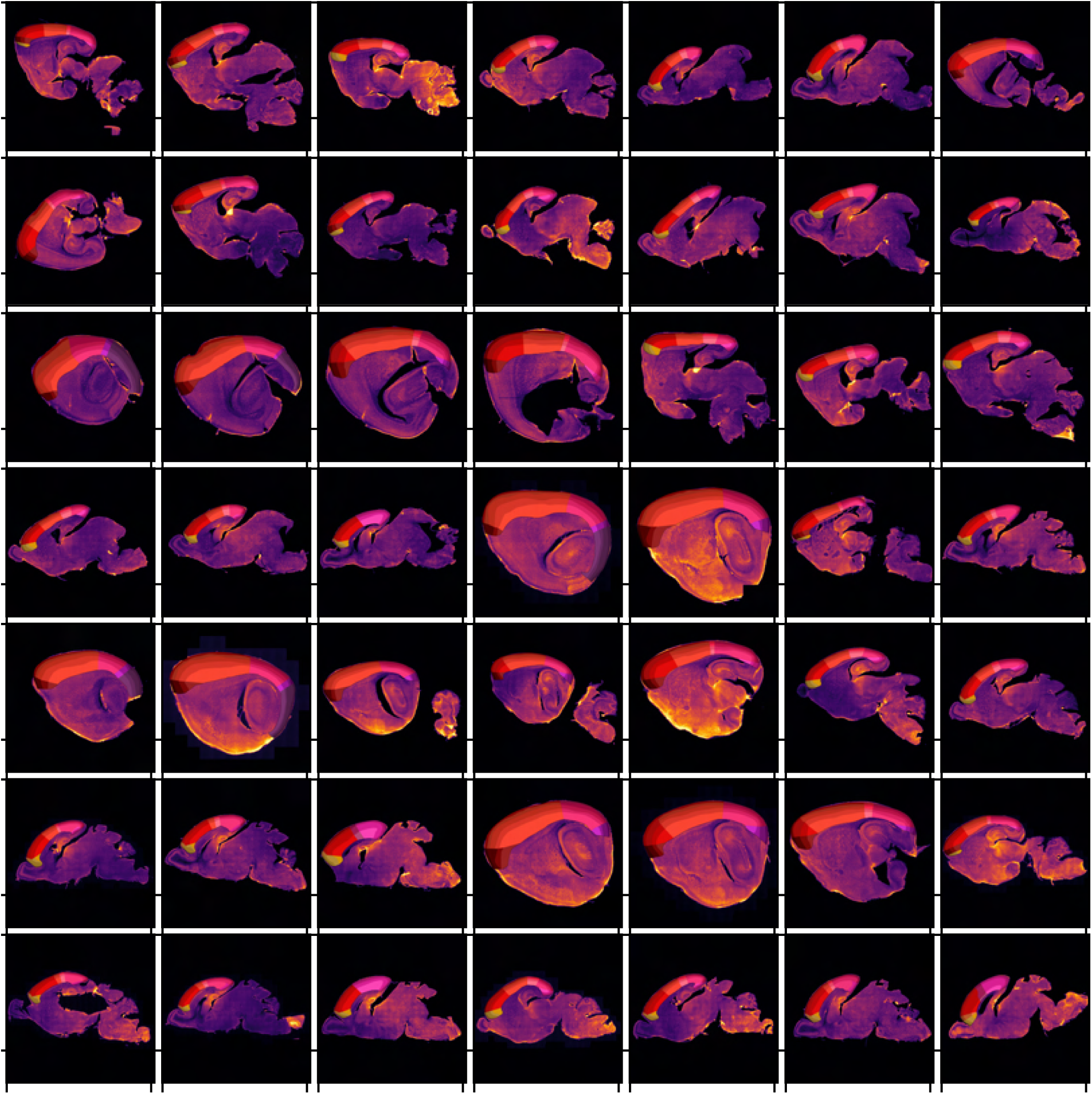
Sample brain sections images from mouse age P4, labeled using the fluorescent pan-neuronal marker, NeuN. All brain sections are converted to grayscale and visualised in the inferno colormap, to allow comparisons between the distinct spatial expression patterns of each brain marker. All brain sections are overlaid with the corresponding manually-annotated cortical atlas, as provided in the MortX dataset.

**Supplementary Figure 11:**
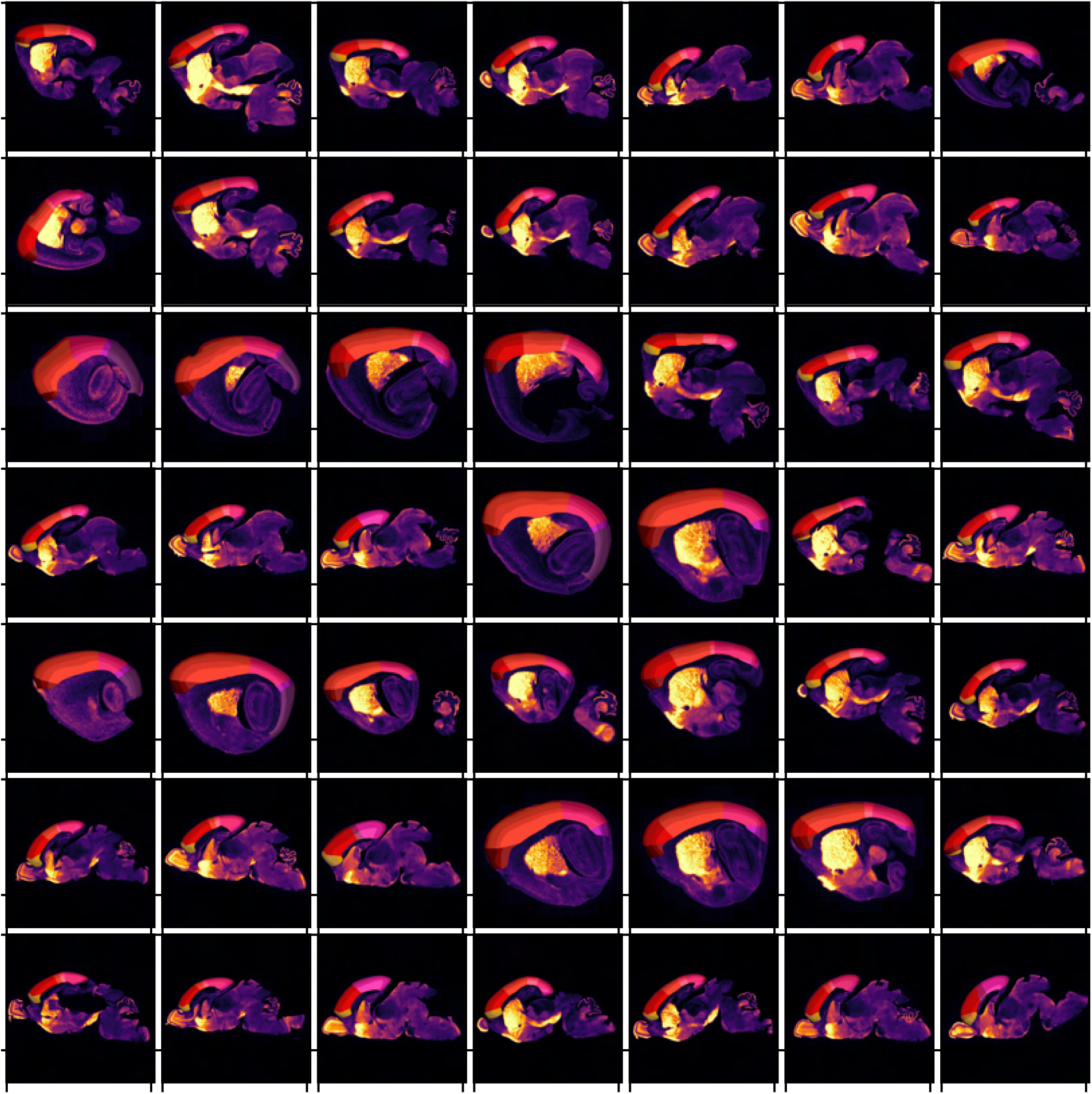
Sample brain sections images from mouse age P4, labelled using the fluorescent inhibitory neuron marker, GAD2. All brain sections are converted to grayscale and visualised in the inferno colormap, to highlight the distinct spatial expression patterns of each brain marker. All brain sections are overlaid with the corresponding manually-annotated cortical atlas, as provided in the MortX dataset.

**Supplementary Figure 12:**
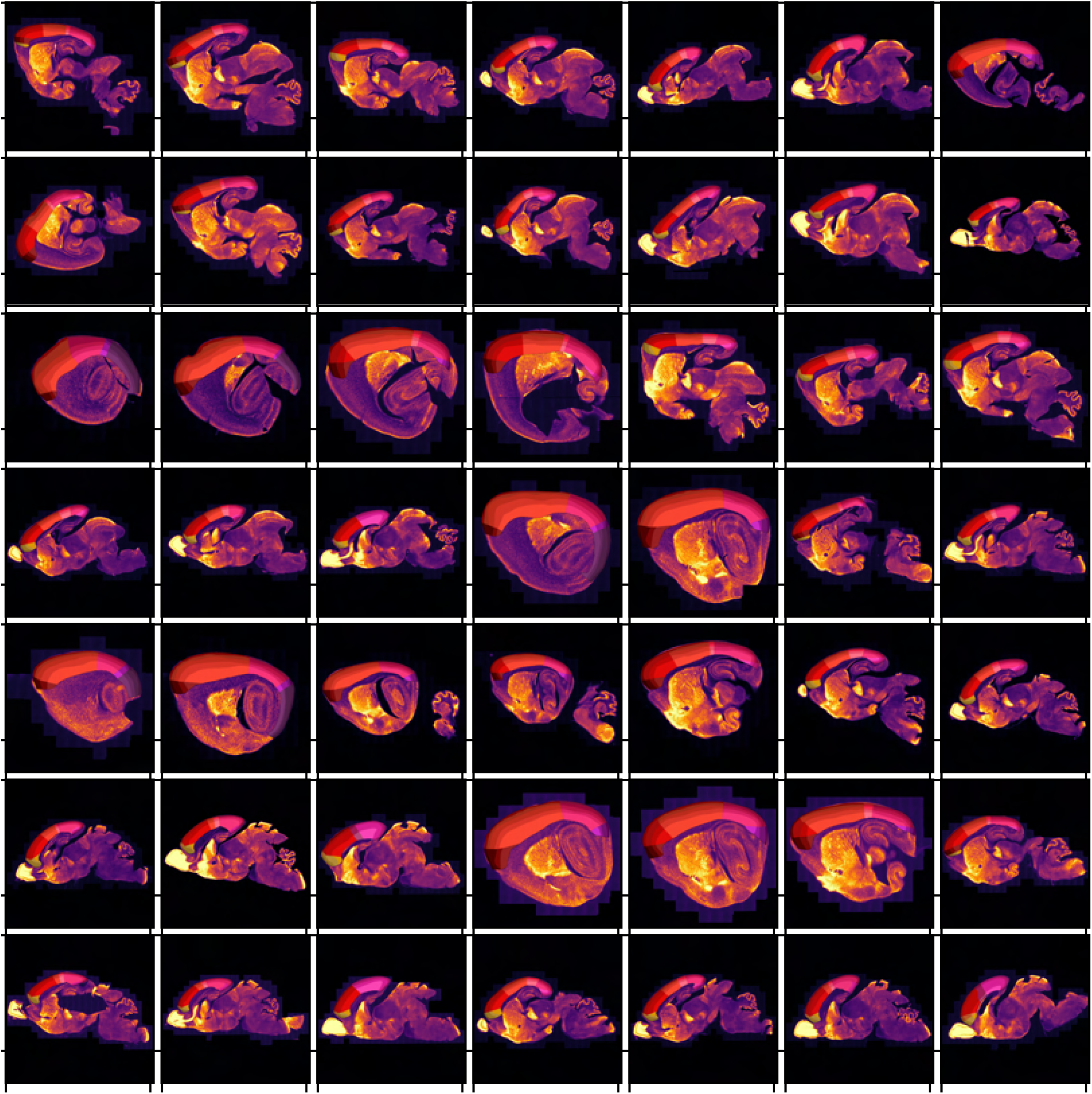
Sample brain sections images from mouse age P4, labelled using the fluorescent inhibitory neuron marker, GAD1. All brain sections are converted to grayscale and visualised in the inferno colormap, to highlight the distinct spatial expression patterns of each brain marker. All brain sections are overlaid with the corresponding manually-annotated cortical atlas, as provided in the MortX dataset.

**Supplementary Figure 13:**
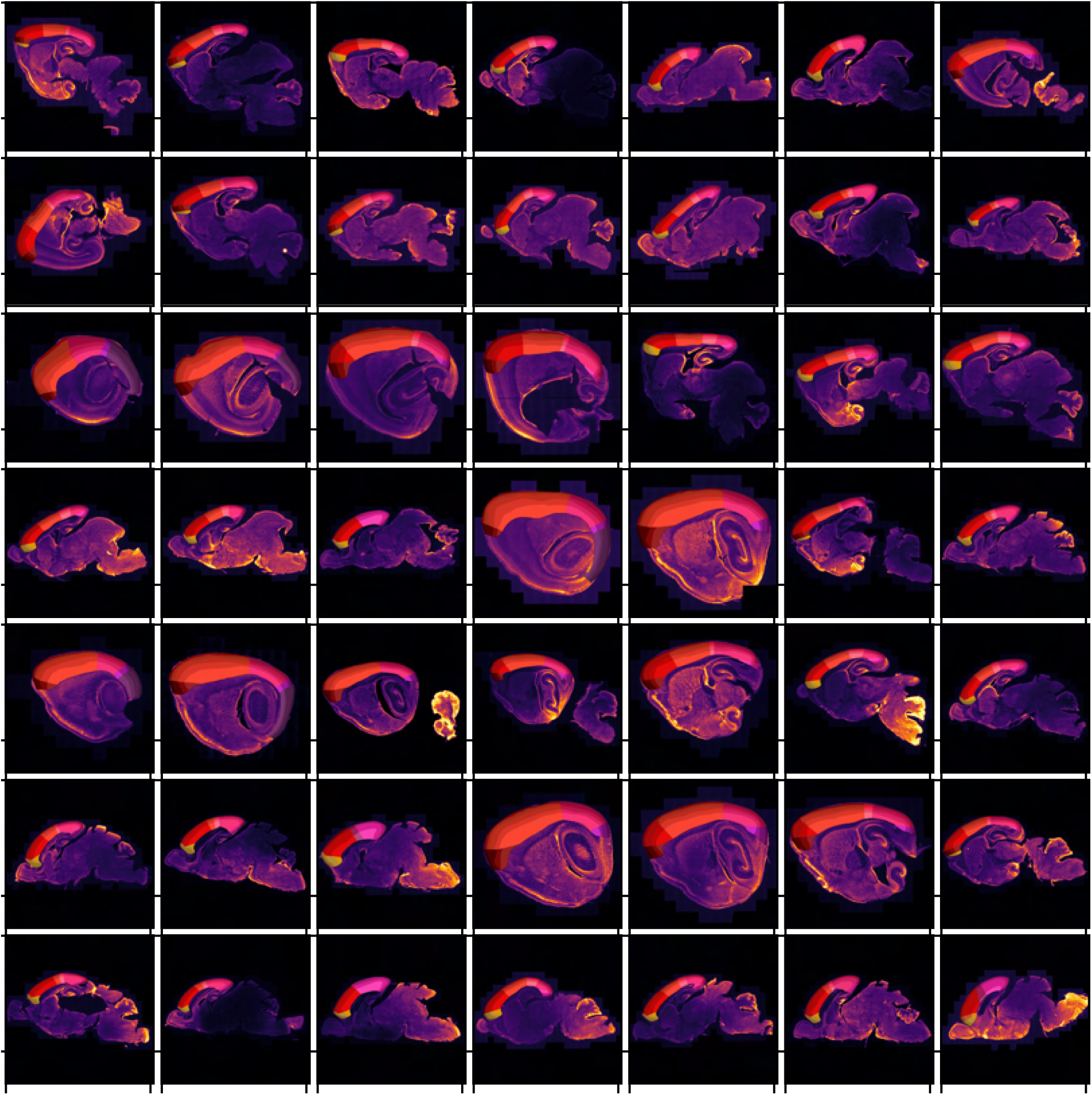
Sample brain sections images from mouse age P4, labeled using the pan-cellular marker, DAPI. All brain sections are converted to grayscale and visualised in the inferno colormap, to highlight the distinct spatial expression patterns of each brain marker. All brain sections are overlaid with the corresponding manually-annotated cortical atlas, as provided in the MortX dataset.

**Supplementary Figure 14:**
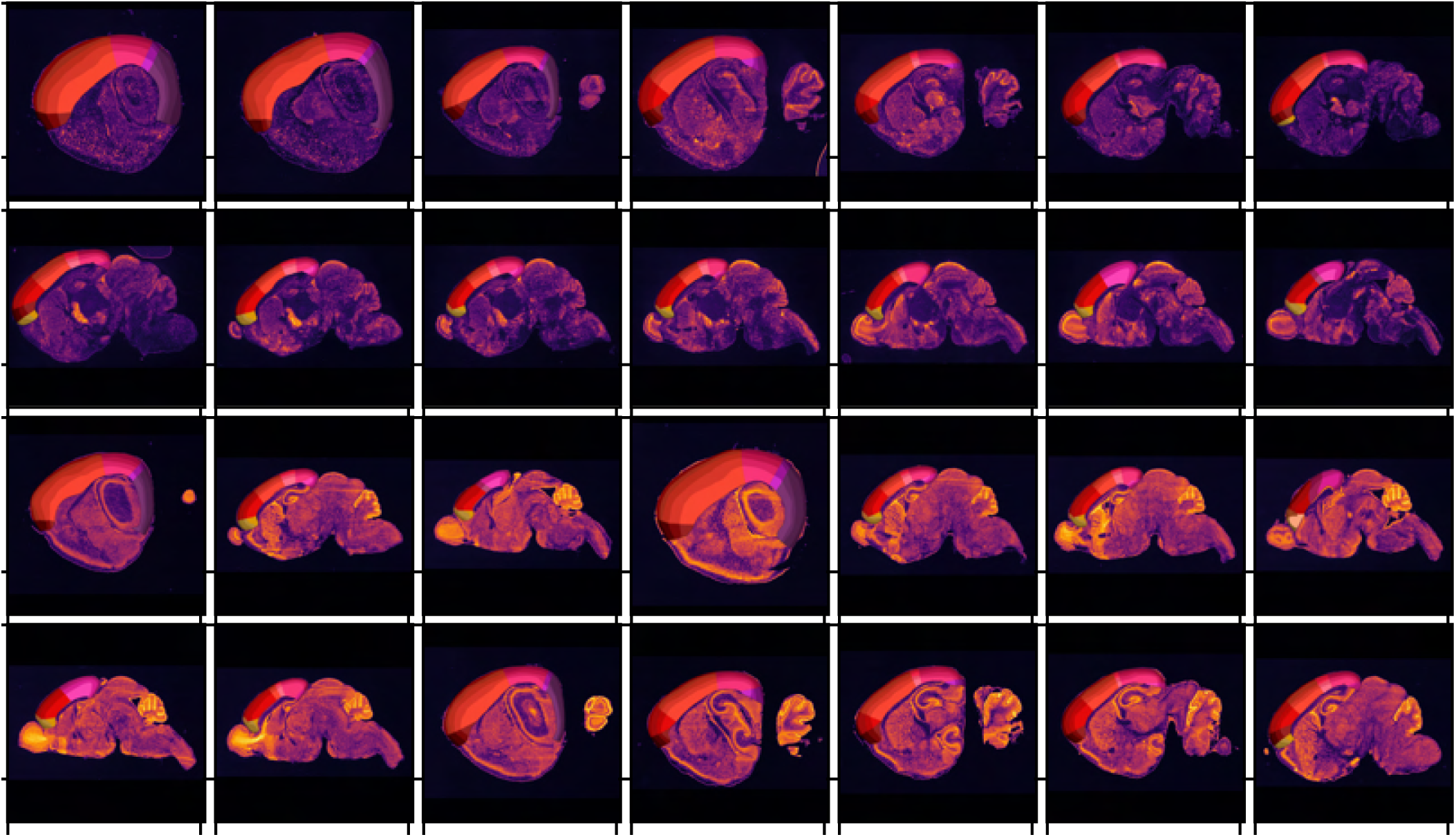
Sample brain sections images from mouse age P4, labelled using the inhibitory neuron marker, GAD1 (rows 3-4), and the nuclear stain, Nissl (rows 1-2), visualized using bright-field imaging. All brain sections are converted to grayscale and visualised in the inferno colormap, to highlight the distinct spatial expression patterns of each brain marker. All brain sections are overlaid with the corresponding manually-annotated cortical atlas, as provided in the MortX dataset.

**Supplementary Figure 15:**
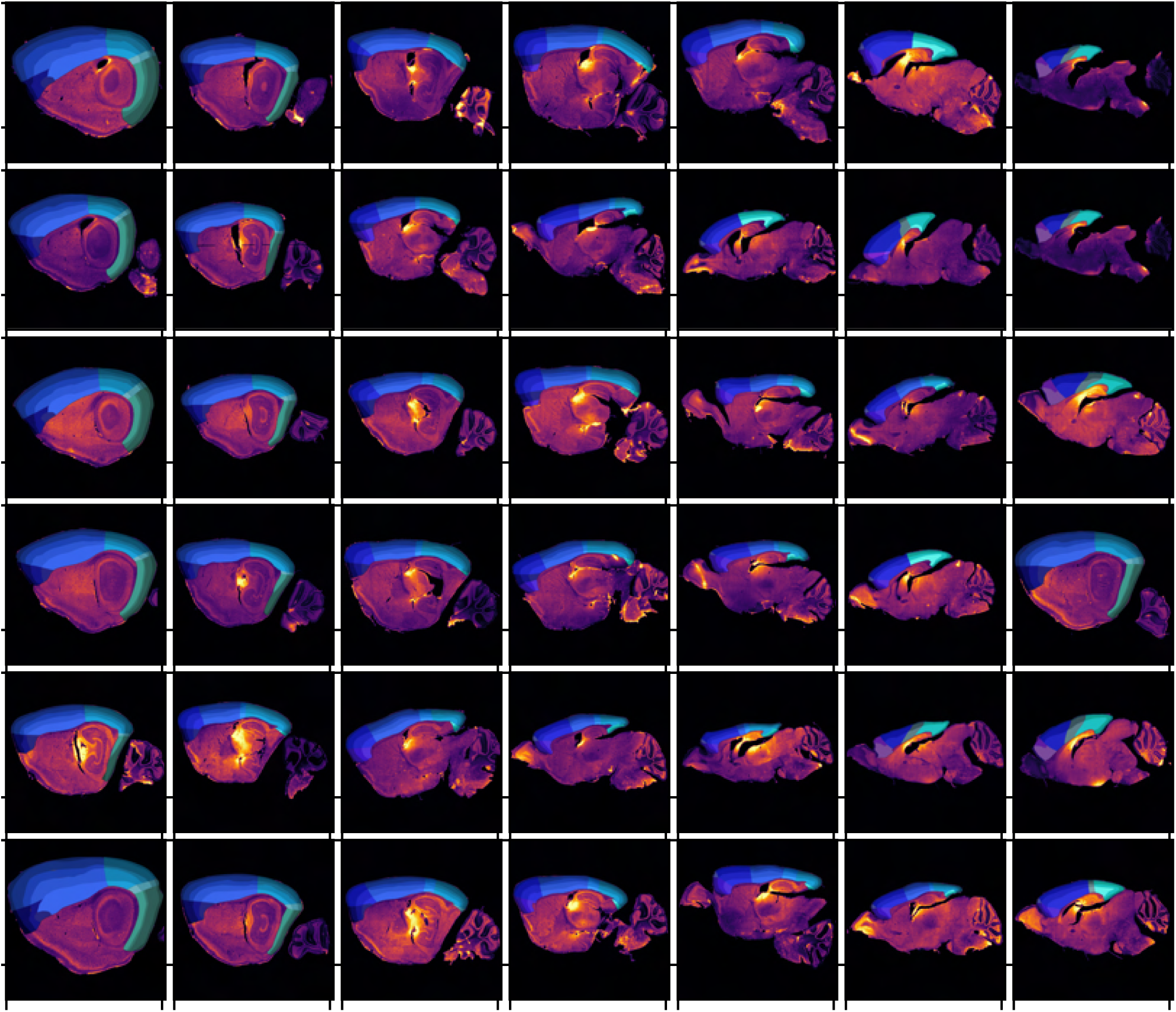
Sample brain sections images from mouse age P14, labeled using the fluorescent pan-neuronal marker, NeuN. All brain sections are converted to grayscale and visualised in the inferno colormap, to allow comparisons between the distinct spatial expression patterns of each brain marker. All brain sections are overlaid with the corresponding manually-annotated cortical atlas, as provided in the MortX dataset.

**Supplementary Figure 16:**
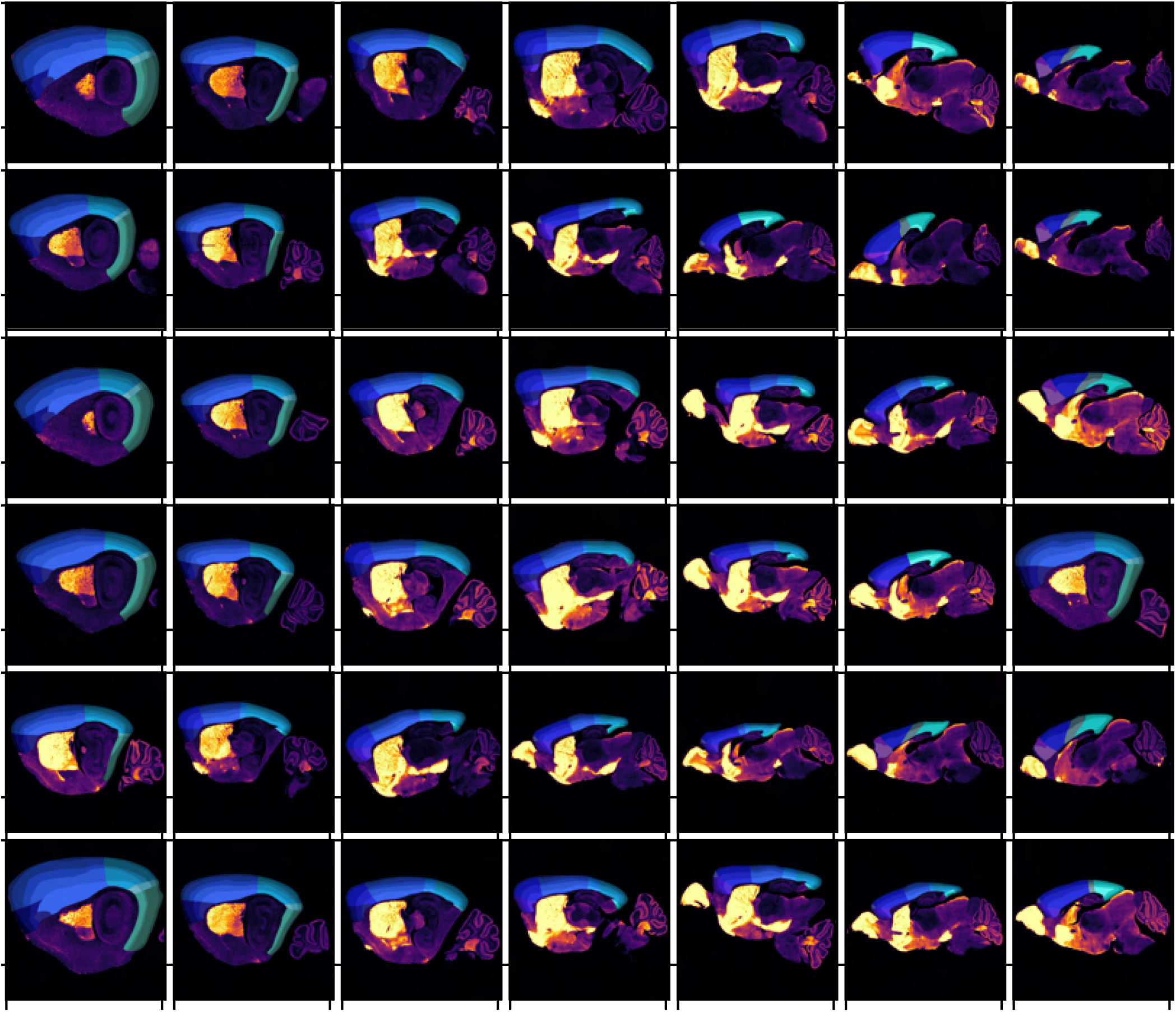
Sample brain sections images from mouse age P14, labelled using the fluorescent inhibitory neuron marker, GAD2. All brain sections are converted to grayscale and visualised in the inferno colormap, to highlight the distinct spatial expression patterns of each brain marker. All brain sections are overlaid with the corresponding manually-annotated cortical atlas, as provided in the MortX dataset.

**Supplementary Figure 17:**
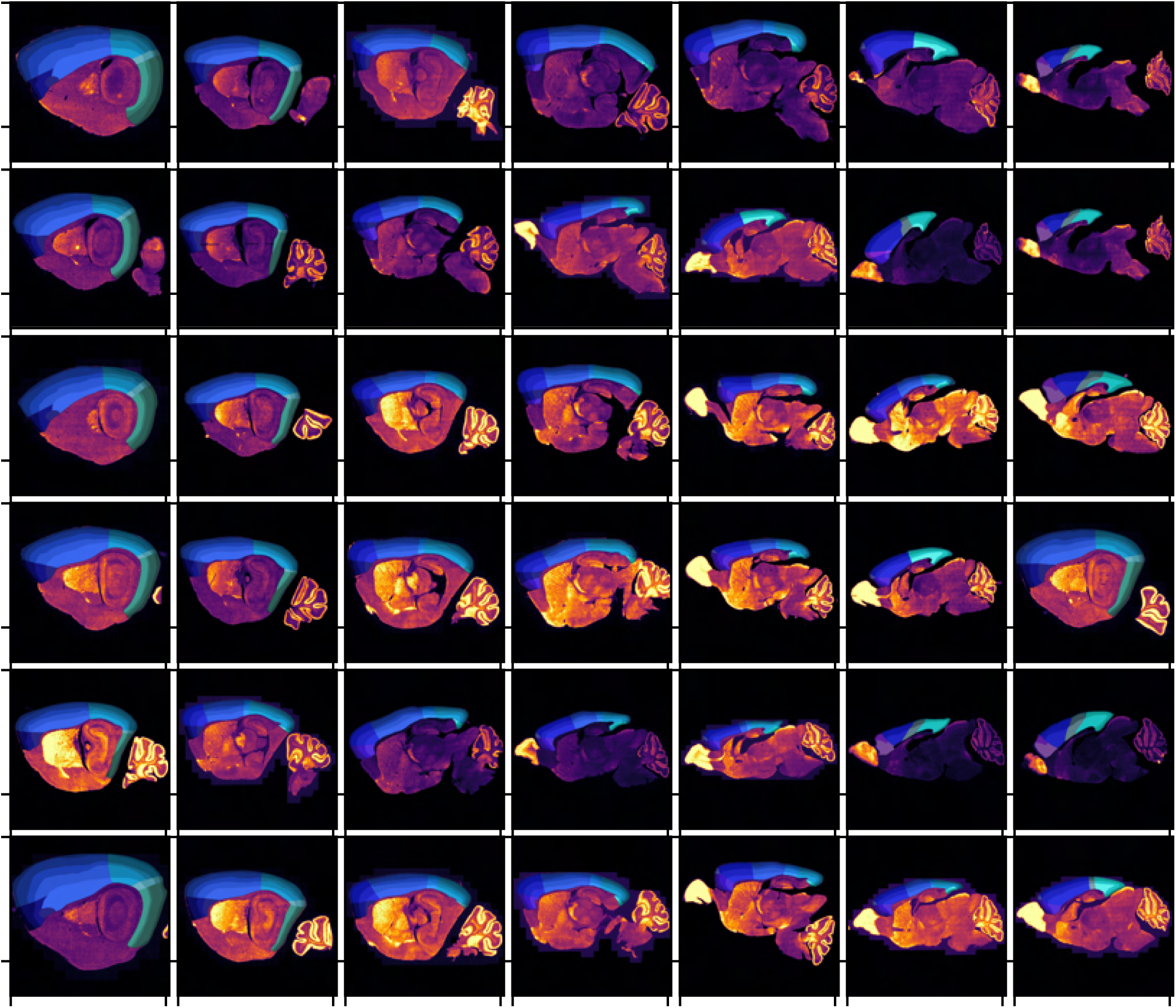
Sample brain sections images from mouse age P14, labelled using the fluorescent inhibitory neuron marker, GAD1. All brain sections are converted to grayscale and visualised in the inferno colormap, to highlight the distinct spatial expression patterns of each brain marker. All brain sections are overlaid with the corresponding manually-annotated cortical atlas, as provided in the MortX dataset.

**Supplementary Figure 18:**
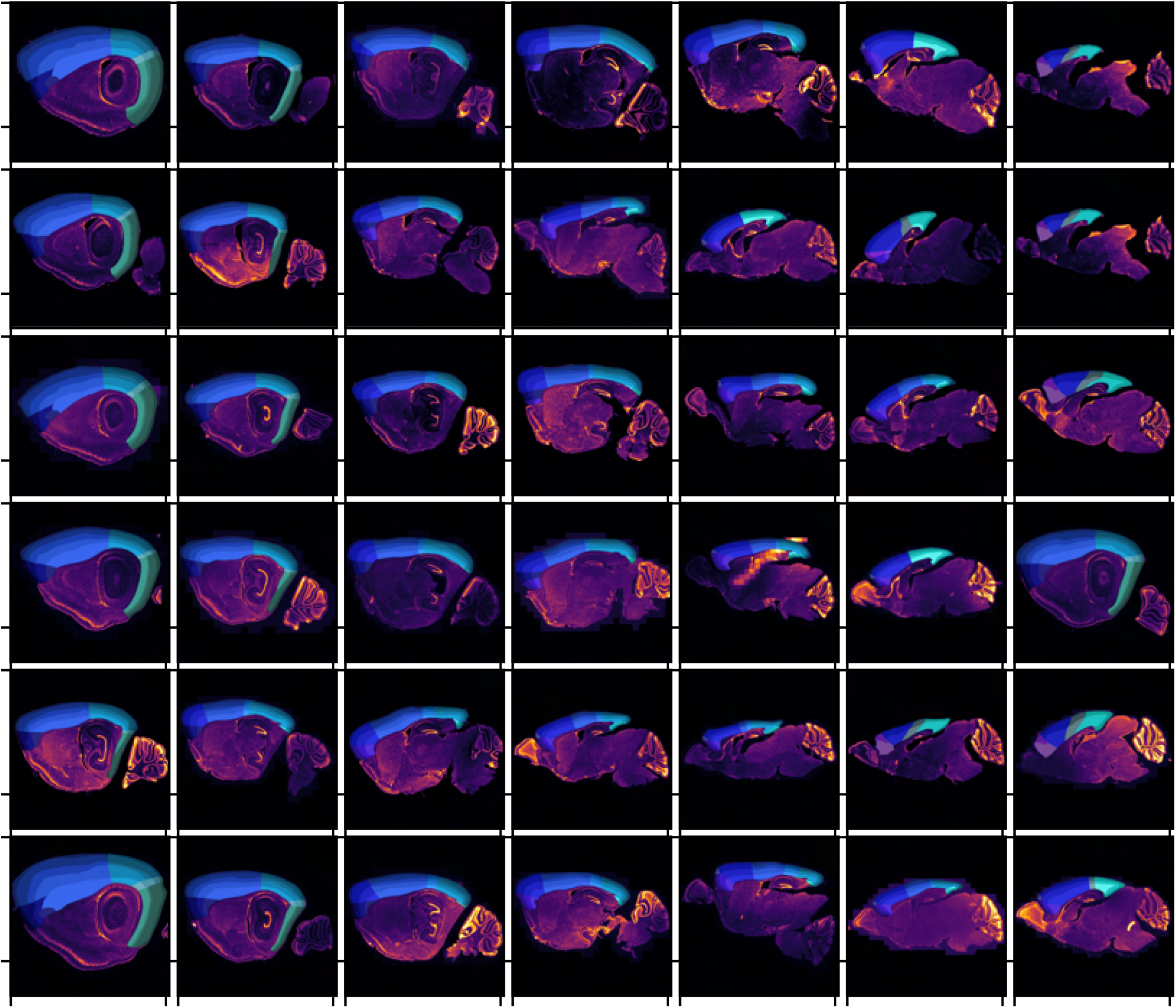
Sample brain sections images from mouse age P14, labeled using the pan-cellular marker, DAPI. All brain sections are converted to grayscale and visualised in the inferno colormap, to highlight the distinct spatial expression patterns of each brain marker. All brain sections are overlaid with the corresponding manually-annotated cortical atlas, as provided in the MortX dataset.

**Supplementary Figure 19:**
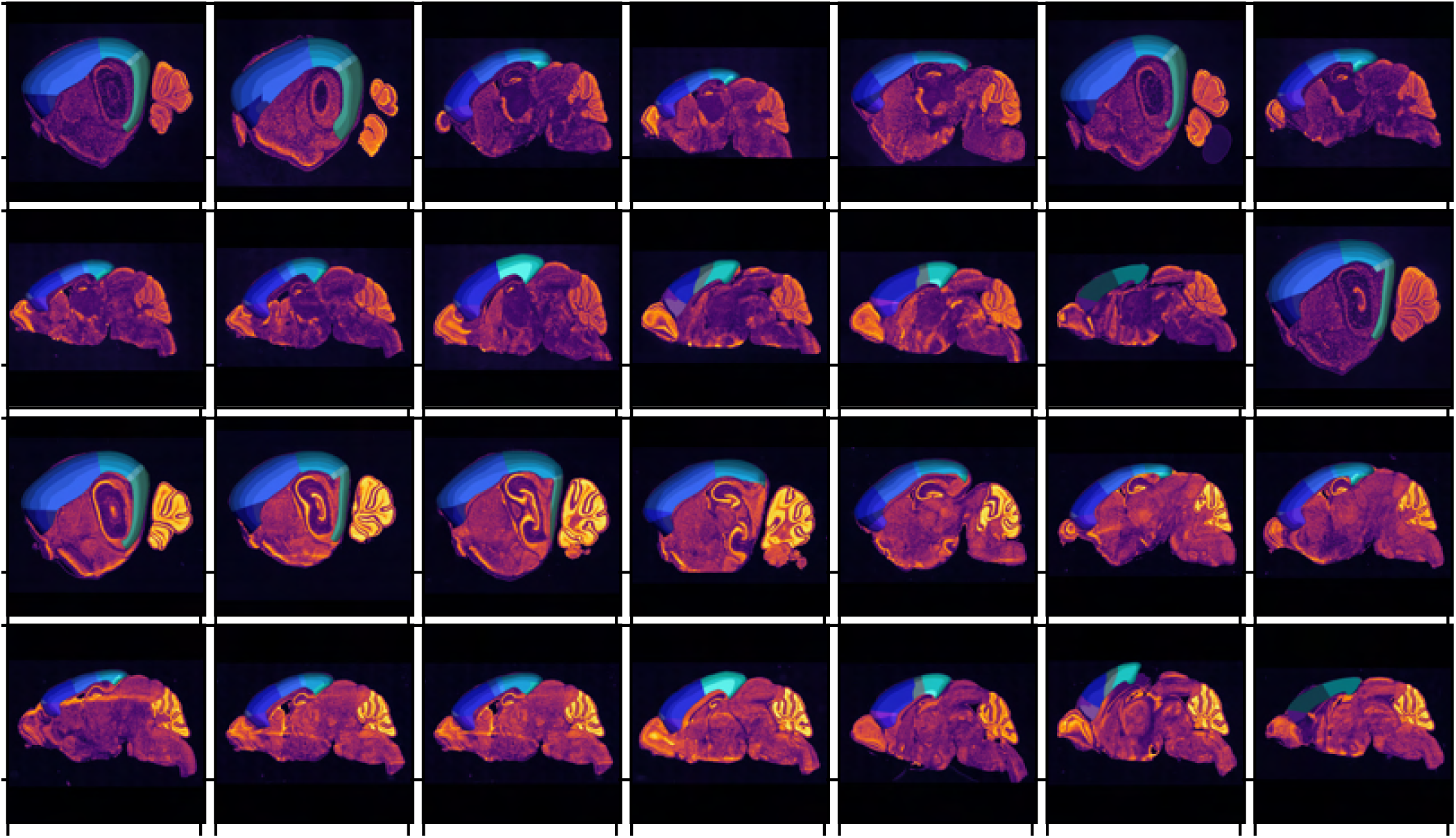
Sample brain sections images from mouse age P14, labelled using the inhibitory neuron marker, GAD1 (rows 3-4), and the nuclear stain, Nissl (rows 1-2), visualized using bright-field imaging. All brain sections are converted to grayscale and visualised in the inferno colormap, to highlight the distinct spatial expression patterns of each brain marker. All brain sections are overlaid with the corresponding manually-annotated cortical atlas, as provided in the MortX dataset.

**Supplementary Figure 20:**
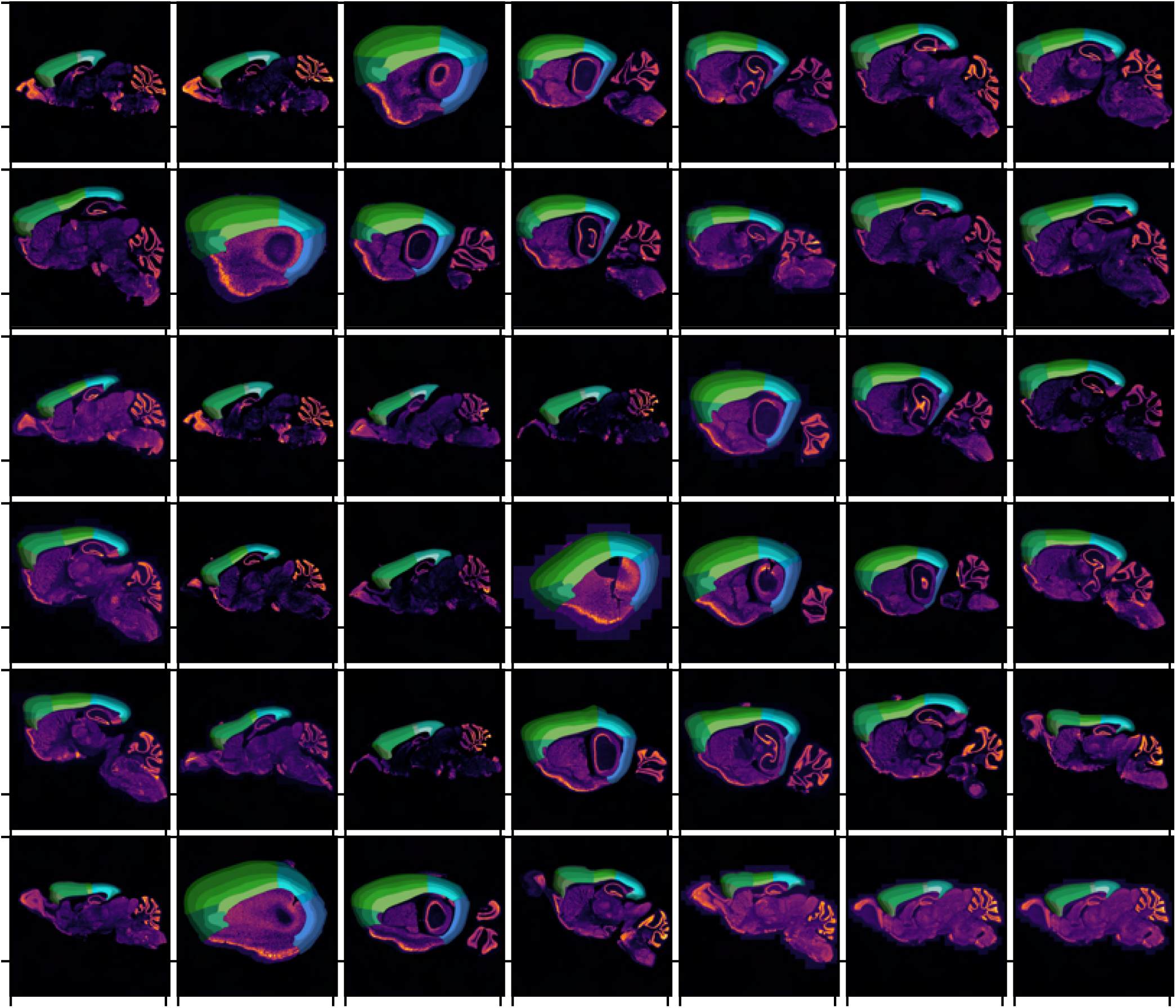
Sample brain sections images from mouse age P56, labeled using the fluorescent pan-neuronal marker, NeuN. All brain sections are converted to grayscale and visualised in the inferno colormap, to allow comparisons between the distinct spatial expression patterns of each brain marker. All brain sections are overlaid with the corresponding manually-annotated cortical atlas, as provided in the MortX dataset.

**Supplementary Figure 21:**
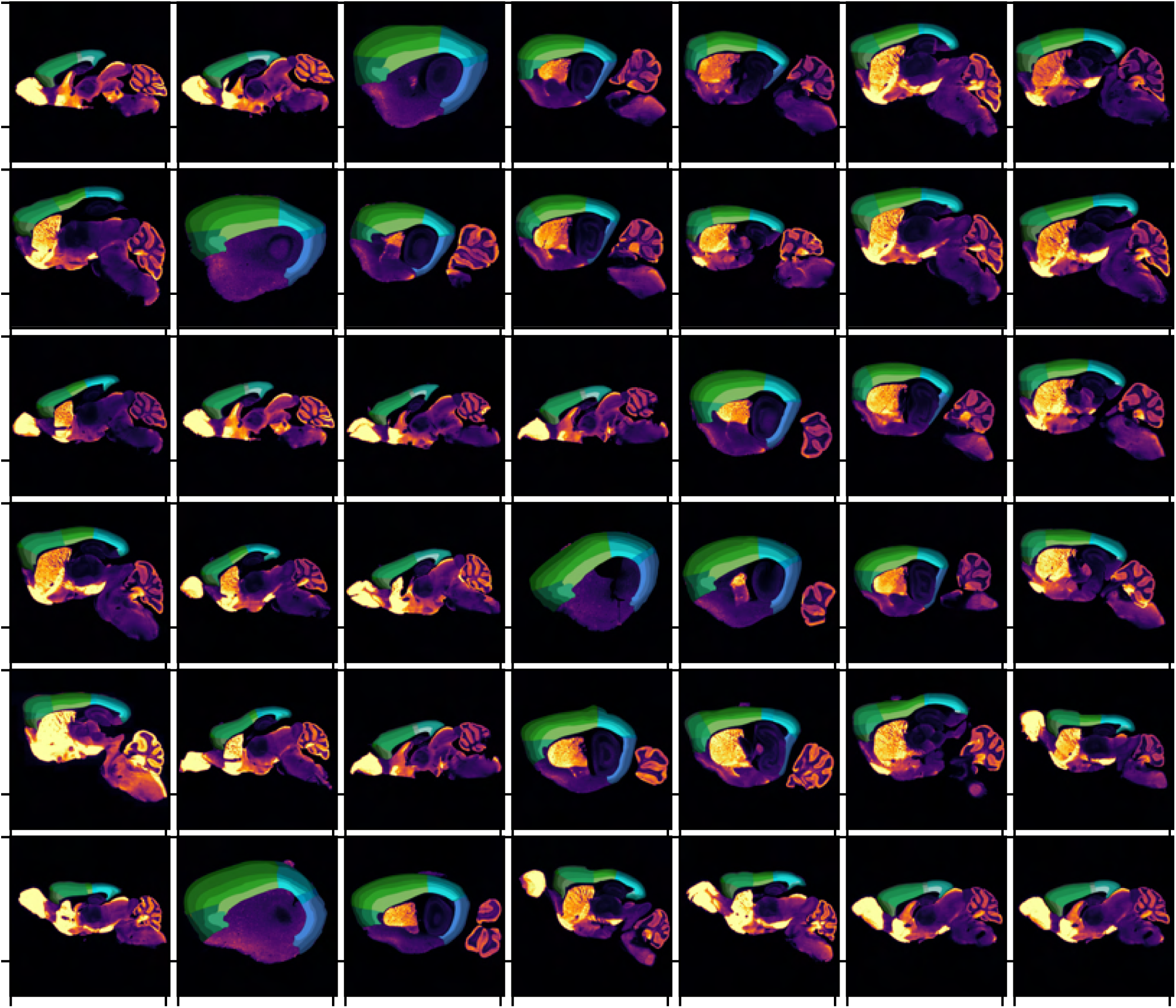
Sample brain sections images from mouse age P56, labelled using the fluorescent inhibitory neuron marker, GAD2. All brain sections are converted to grayscale and visualised in the inferno colormap, to highlight the distinct spatial expression patterns of each brain marker. All brain sections are overlaid with the corresponding manually-annotated cortical atlas, as provided in the MortX dataset.

**Supplementary Figure 22:**
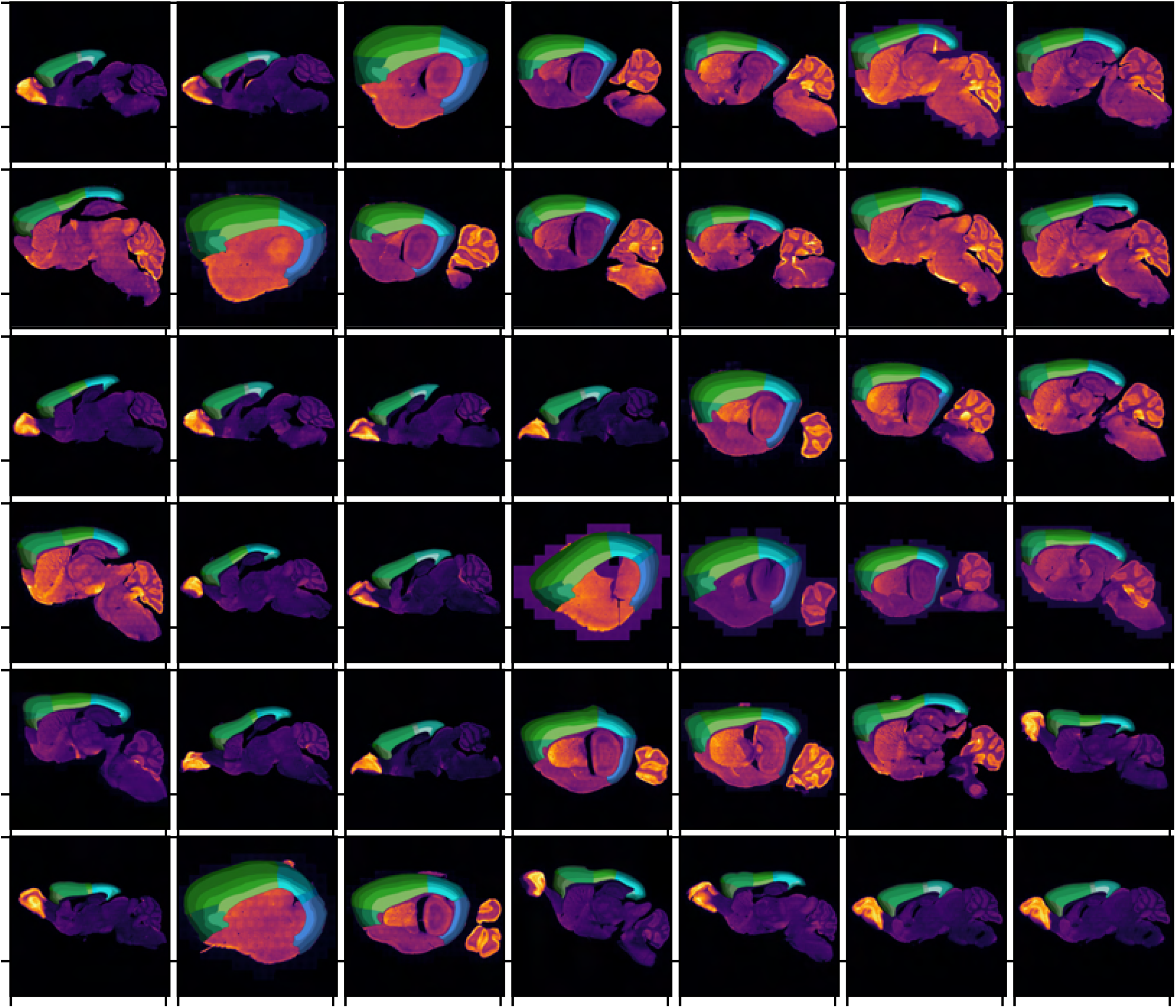
Sample brain sections images from mouse age P56, labelled using the fluorescent inhibitory neuron marker, GAD1. All brain sections are converted to grayscale and visualised in the inferno colormap, to highlight the distinct spatial expression patterns of each brain marker. All brain sections are overlaid with the corresponding manually-annotated cortical atlas, as provided in the MortX dataset.

**Supplementary Figure 23:**
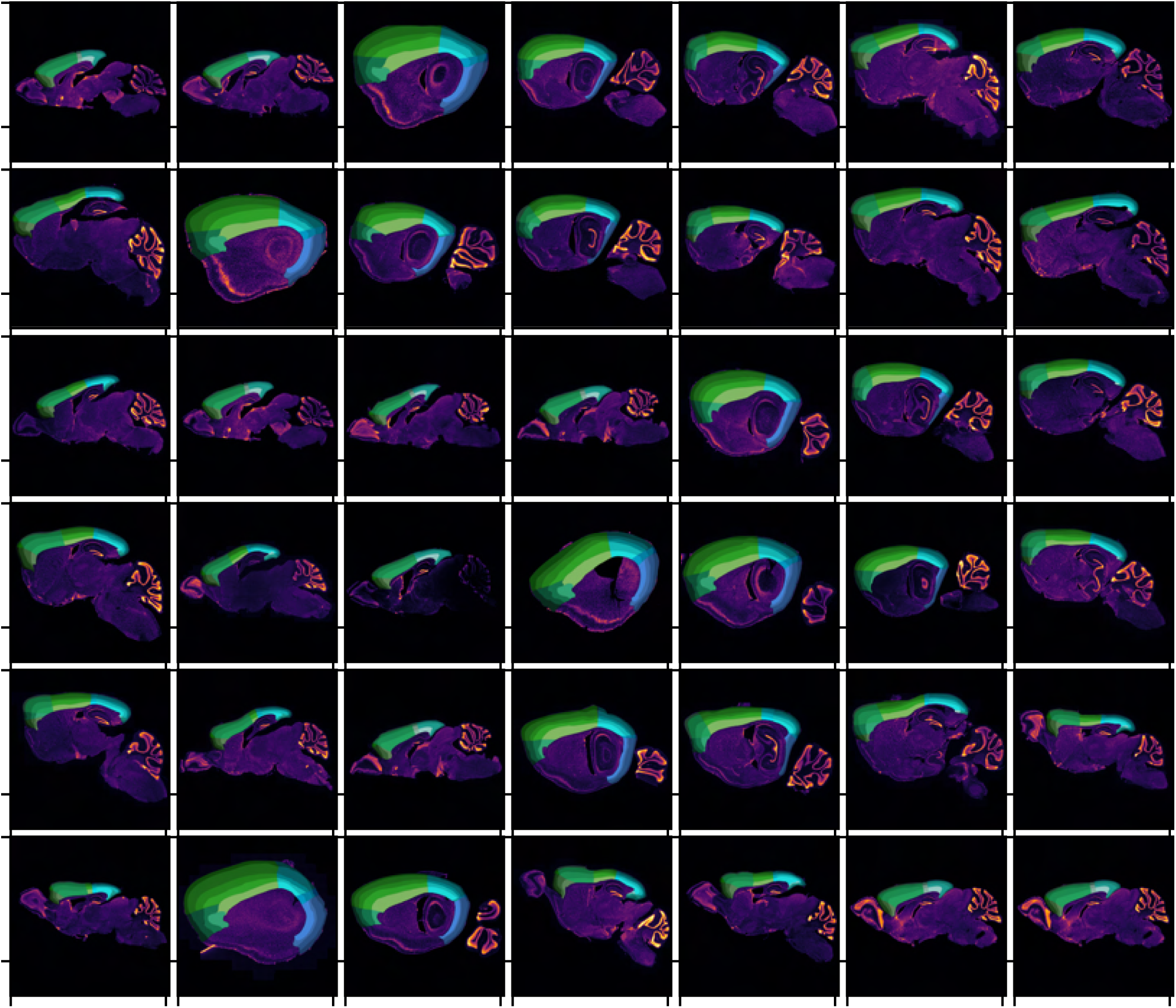
Sample brain sections images from mouse age P56, labeled using the pan-cellular marker, DAPI. All brain sections are converted to grayscale and visualised in the inferno colormap, to highlight the distinct spatial expression patterns of each brain marker. All brain sections are overlaid with the corresponding manually-annotated cortical atlas, as provided in the MortX dataset.

**Supplementary Figure 24:**
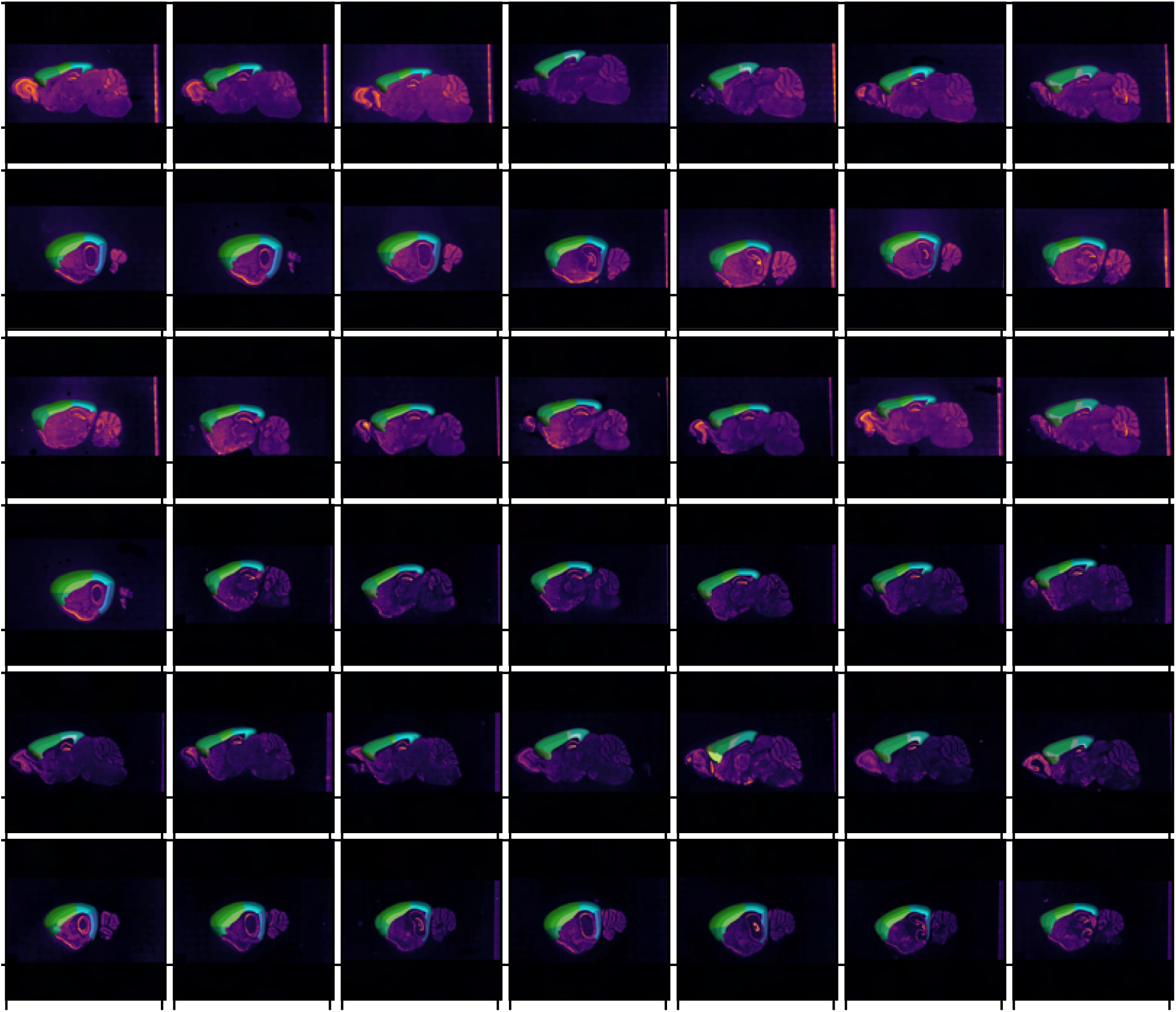
Sample brain sections images from mouse age P56, co-labelled using the inhibitory neuron marker, GAD1, and the excitatory marker, CamK2, visualized using bright-field imaging. All brain sections are converted to grayscale and visualised in the inferno colormap, to highlight the distinct spatial expression patterns of each brain marker. All brain sections are overlaid with the corresponding manually-annotated cortical atlas, as provided in the MortX dataset.

**Supplementary Figure 25:**
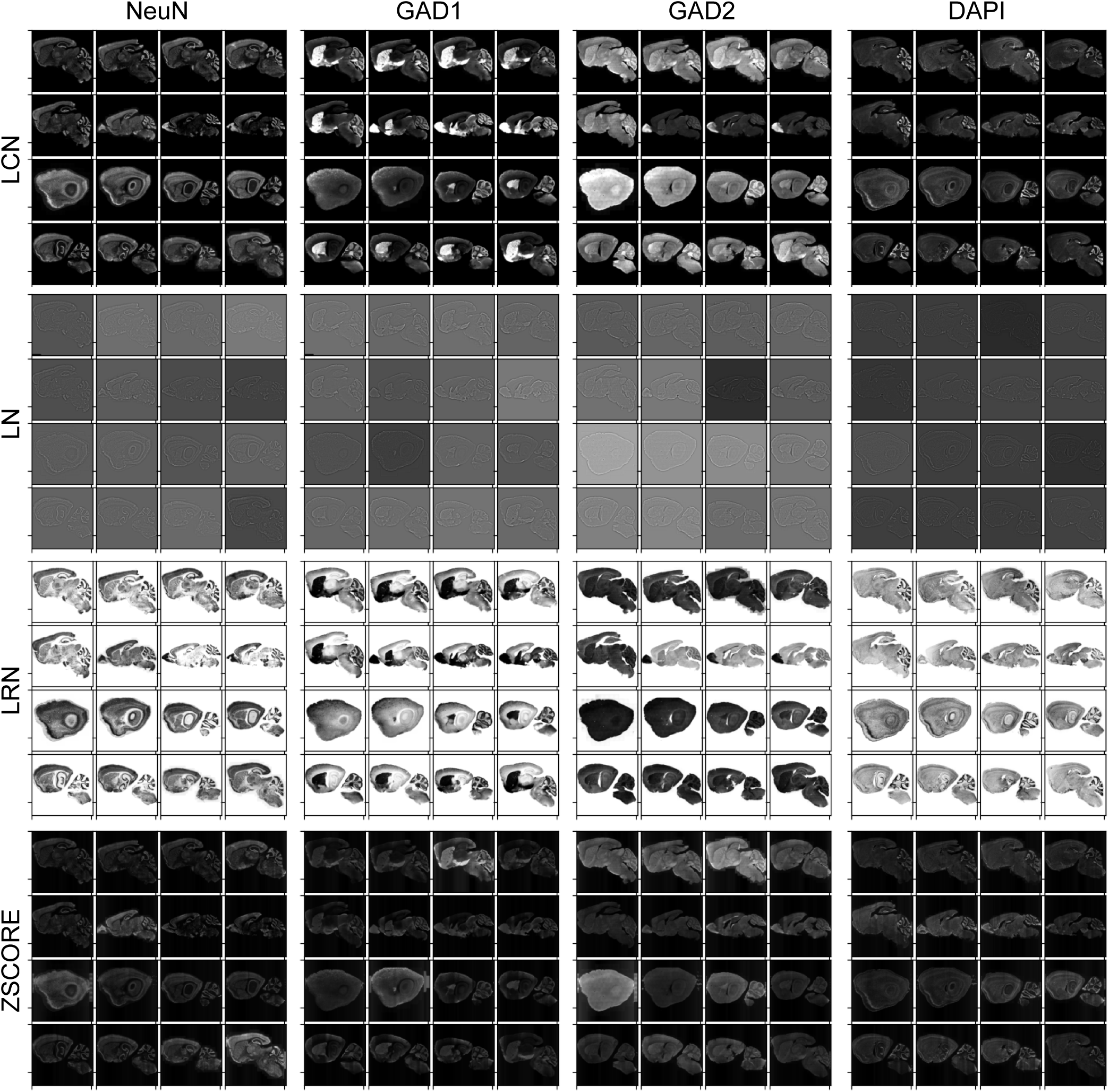
Brain section images labelled with different brain markers, NeuN (column 1), GAD1 (column 2), GAD1 (column 3) and DAPI (column 4), are visualised after preprocessing using common domain adaptation methods: LCN (row 1), LN (row 2), LRN (row 3), and z-score normalization (row 4).

**Supplementary Figure 26:**
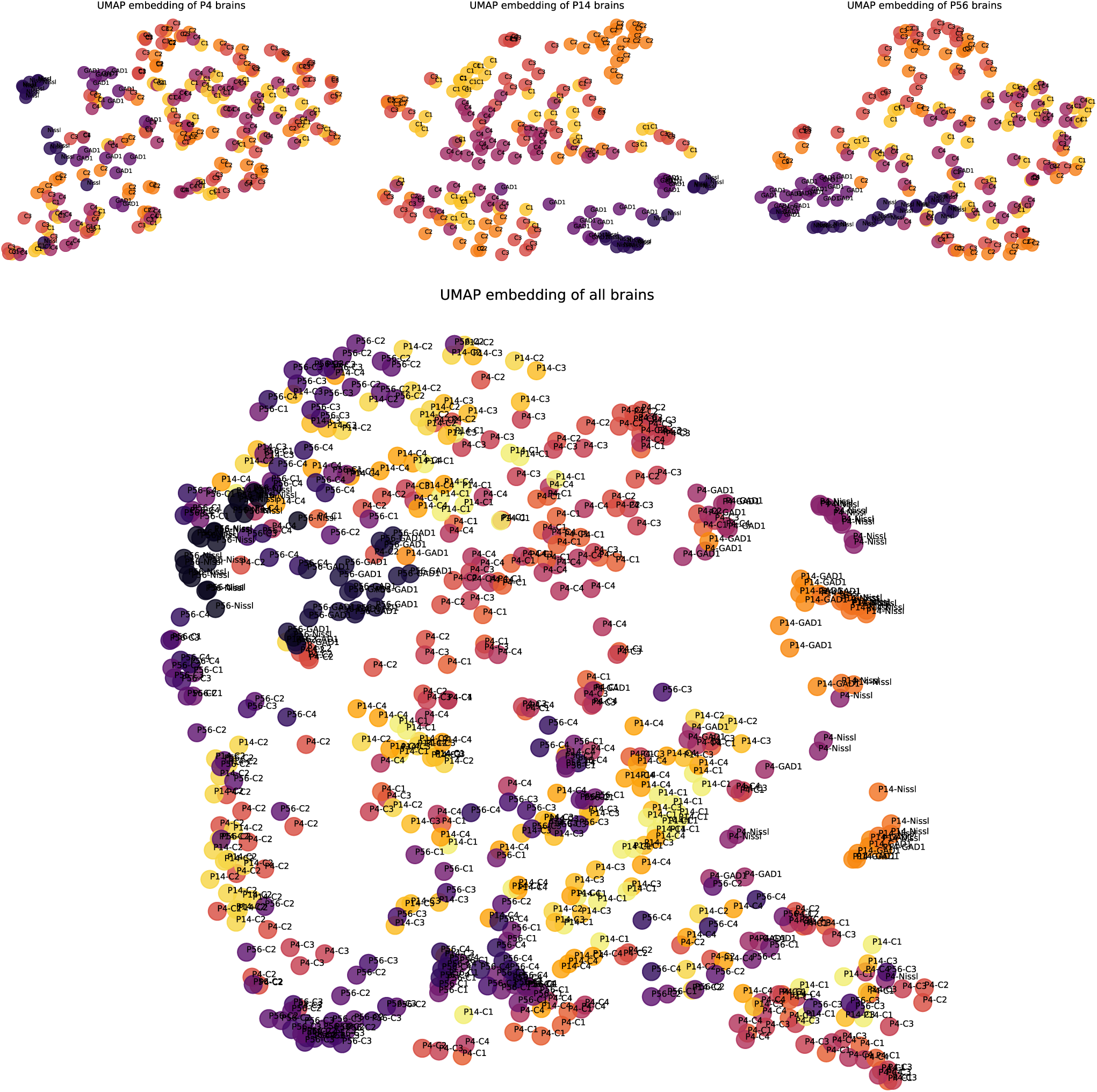
All raw brain section images are represented as 2-dimensional UMAP embeddings, plotted as scatterplots for each mouse age (P4, P14, and P56), in the top panel. In the bottom panel, the embeddings corresponding to all brain sections in the dataset are presented together. Each color code represents a unique genetically-labelled mouse brain. C1: NeuN, C2: GAD2, C3:GAD1, C4:DAPI.

